# Inactivation of microtubule organizing center function of the centrosome is required for neuronal development and centriole elimination

**DOI:** 10.1101/2025.11.17.688953

**Authors:** Rachel Ng, Jérémy Magescas, Jessica L. Feldman

## Abstract

Cell differentiation is marked by a dramatic reorganization of the microtubule cytoskeleton that enables diverse cell functions. Mitotic precursor cells arrange microtubules around centrosomes, which become activated as microtubule-organizing centers (MTOCs) through the recruitment of pericentriolar material (PCM) and microtubules to build the mitotic spindle. A hallmark of differentiation is the “inactivation” of centrosomal MTOC function through the loss of PCM and microtubules, yet the function of this inactivation is unknown. We developed a GFP-nanobody-based targeting tool to activate centrosomes in *C. elegans* differentiated cells. Ectopically activating centrosomes in sensory neurons perturbed microtubule polarity, dynein-mediated trafficking, and caused defects in cell morphology and dendrite pathfinding. Ectopic PCM perturbed ciliogenesis and also protected centrioles from elimination, another common feature of differentiation. By forcing centrosome activation in a fully differentiated cell in a developing organism, we show that centrosome inactivation is required for differentiation by directly contributing to cell form and function.

## Introduction

Diverse microtubule arrays support specialized cell forms and functions and are generated by microtubule organizing centers (MTOCs) that nucleate and organize microtubules into cell-type and cell-state specific patterns. In animal mitosis, organelles called centrosomes function as MTOCs to help build the mitotic spindle. As cells differentiate, however, centrosomes are often “inactivated” as MTOCs, and other subcellular sites act as non-centrosomal MTOCs (ncMTOCs) to generate diverse microtubule arrays ^1^. For example, in epithelial cells, the apical surface and/or cell junctions generate parallel microtubule arrays along their apical-basal axis ^2–4^. In striated muscle cells, the nuclear envelope acts as an MTOC to generate large, radial microtubule arrays ^5^. In neurons, axons and dendrites contain linear arrays of microtubules with specific polarity: axons across organisms universally contain microtubules with their plus ends pointing away from the cell body, while dendrites can have some or all of their microtubules oriented with plus ends pointing towards the cell body ^6,7^ that can be generated by ncMTOCs found on endosomes or Golgi-derived structures ^8–10^. This microtubule polarity can also be reinforced through motor-based microtubule sliding ^11–15^. Thus, while the establishment of ncMTOCs in differentiated cells is well documented, whether the inactivation of MTOC function at the centrosome is a necessary step in differentiation has never been tested.

Centrosomes are membrane-less organelles composed of a pair of microtubule-based centrioles surrounded by proteins known collectively as the pericentriolar material (PCM). In mitosis in animal cells, centrosomes become activated as MTOCs through the kinase-dependent recruitment of PCM and the ɣ-tubulin ring complex (ɣ-TURC) around the centrioles to nucleate the microtubules of the mitotic spindle ^16^. In telophase, centrosomes are inactivated or severely attenuated as MTOCs through the loss of PCM proteins, which is in part controlled by phosphatase activity ^17,18^. Apart from the maintenance of this inactive state as cells differentiate, centrosomes can also be terminally inactivated through the elimination of the centrioles such as in cardiomyocytes and other striated muscle cells, oocytes, enterocytes, and the majority of cells in *C. elegans* ^19–22^. Thus, centrosome inactivation is a ubiquitous process that occurs at the end of every cell division and during differentiation, but the function of centrosome inactivation is unknown. Correlative studies have also shown that hyperactive centrosomal MTOC activity is a hallmark of tumors and has been linked to many cancers and invasive cell behaviors ^23,24^. Thus, the inactivation of MTOC activity at the centrosome may not only be a key step in cell differentiation, but might also help suppress pathogenesis, although direct tests of the role of centrosome inactivation in differentiated cells need to be performed.

While the mechanisms underlying centrosome inactivation in differentiated cells are relatively unknown, phosphatase activity and microtubule-based mechanical forces drive the loss of PCM at the end of mitosis and the precocious loss of CDK1 activity causes premature PCM disassembly ^17,18,25^. This disassembly process removes mitotic PCM, likely leaving a small shell of interphase PCM in most cell types. The complete loss of PCM has been suggested to be an important step prior to centriole elimination during female gametogenesis, with ectopic centrosomal MTOC activity protecting centrioles from elimination and interfering with female meiotic spindle assembly ^21^.

Similarly, the MTOC activity of the first centrosome brought into the zygote from the sperm during fertilization must be suppressed, in part through kinesin I activity ^26^.

Indeed, precociously active sperm centrosomes interfere with female meiosis and subsequent cell division in the embryo ^26^. Despite these limited studies in gametes and decades of observational data demonstrating the inactivation of centrosomal MTOC function during cell differentiation, functional roles for centrosome inactivation, especially *in vivo*, still need to be established.

Studying the function of centrosome inactivation directly has been difficult due to mitotic defects that arise upon perturbing the centrosome and the challenge of uncoupling mitotic defects from phenotypes associated with improper centrosome inactivation. Here, we developed a novel genetic tool to ectopically activate the MTOC function of the centrosome during neuronal differentiation in *C. elegans* while avoiding mitotic defects normally associated with centrosome perturbation. Using this tool, we have, for the first time, activated centrosomes in a fully differentiated non-dividing cell within a developing organism *in vivo*. Focusing on a pair of ciliated sensory neurons in the tail of the worm, we found that hyperactive centrosomal MTOC function perturbed the stereotyped dendrite microtubule polarity and downstream polarized microtubule-based cargo trafficking. Centrosome activation also perturbed dendrite morphology through aberrant microtubule nucleation and ciliary transition-zone defects. Ectopic PCM recruited to active centrosomes perturbed ciliogenesis and also prevented normal centriole elimination, likely protecting the centriole from degradation machinery in the cytoplasm. These studies reveal for the first time that ectopically activating the MTOC activity of the centrosome in a differentiated cell leads to severe consequences on cell form and function.

## Results

### CAP-Trap activates MTOC function at centrosomes in *C. elegans* phasmid neurons

Centrosomes are activated as MTOCs in mitosis through the phosphorylation of PCM proteins. Mitotic kinases like PLK-1/Polo phosphorylate PCM proteins such as SPD-5, the functional ortholog of Cnn/CDK5RP2, SPD-2/CEP192, and PCMD-1, leading to their multimeric expansion around the centriole and the downstream recruitment of y-TuRC to nucleate microtubules ^27–32^. Due to this ability of PLK-1 to drive PCM expansion and therefore increased MTOC activity, the tethering of PLK-1/Polo to the centrosome had been previously shown to ectopically activate centrosomes in Drosophila oocytes ^21^. We therefore aimed to adapt this strategy for *C. elegans* to test the effect of ectopic centrosome activation in a differentiated cell type. Since PLK-1 is normally activated in mitosis through a priming phosphorylation by the mitotic kinase Aurora A ^33^, we generated a <u>C</u>onstitutively <u>A</u>ctive form of <u>P</u>LK-1 (CAP) to express in interphase by introducing a phospho-mimetic mutation (T194D) at the site of Aurora A priming. To drive CAP to the centriole, we expressed CAP fused to a GFP-nanobody in a *C. elegans* strain in which the centriole protein SAS-7 had been endogenously tagged with GFP (Fig. 1a-b). Together, we refer to this tool as CAP-Trap since the GFP-nanobody will trap CAP at GFP::SAS-7-labeled centrioles (Fig. 1b).

**Figure 1.**
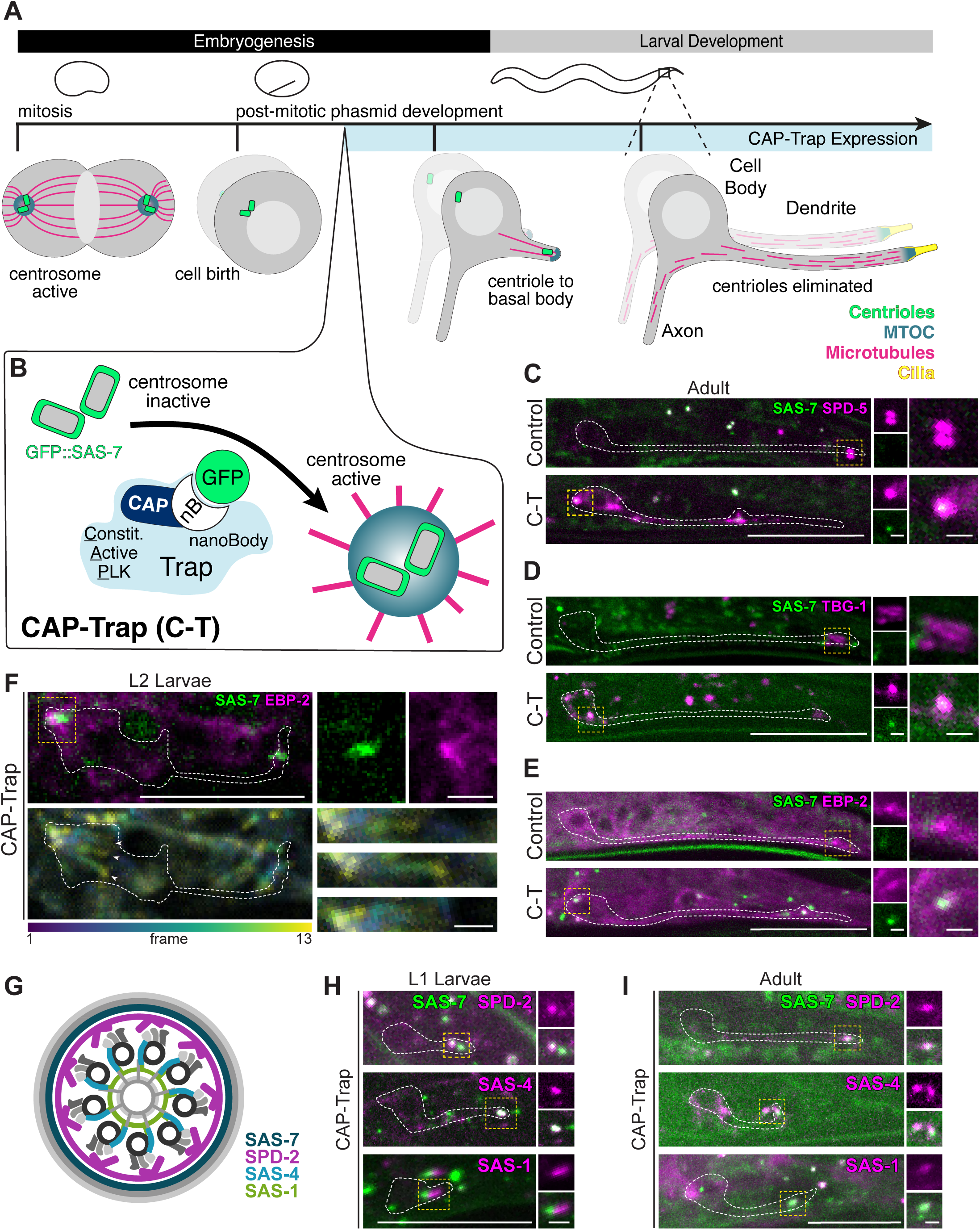
CAP-Trap activates MTOC activity at centrosomes in *C. elegans* phasmid neurons A) Cartoon of phasmid neuron development with centrioles (green), microtubules (magenta), microtubule organizing center (teal), and cilia (yellow) highlighted. B) Schematic of CAP-Trap tool: <u>C</u>onstitutively-<u>A</u>ctive <u>P</u>LK-1(CAP(T194D)) fused to a vhhGFP4-nanobody traps CAP at centrioles through interactions with SAS-7 tagged endogenously with GFP. C-E) Control^No C-T^ (‘No CAP-Trap’) and CAP-Trap (‘C-T’) expressing adult neurons. Magnified insets of boxed regions shown at right. C-E) Control neurons lack GFP::SAS-7 in the dendrite but localize MTOC components to the tip of the dendrite. In contrast, CAP-Trap expressing neurons have SAS-7 puncta that colocalize these components: C) TagRFP::SPD-5 (n=40/40 neuron pairs (NPs)), D) mCherry::TBG-1/ɣ-tubulin (n=32/32 NPs), and E) EBP-2/EB1::mKate2 (n=51/51NPs). In C-T expressing adult neurons, MTOC proteins colocalize with SAS-7 (SPD-5: 91.3% colocalization, n=161 SAS-7 foci, n=55 NPs; TBG-1: 76.1% colocalization, n=67 SAS-7 foci, n=38 NPs; EBP-2: 85.7% colocalization, n=98 SAS-7 foci, n=56 NPs). F) Top: C-T expressing L2 larvae expressing GFP::SAS-7 and EBP-2::mKate-2. Single plane image of EBP-2::mKate2 (magenta) and GFP::SAS-7 (green) with magnified image of boxed region shown at right. Bottom: Time projection of EBP-2 dynamics using the Fiji temporal color code mpl-viridis, 13 frames, 600ms between frames. G) Diagram adapted from Wogler et al, 2022 of the *C. elegans* centriole highlighting proteins localized in H-I. H-I) Colocalization of centriolar proteins with SAS-7 in C-T expressing L1 larvae (H) or adult (I) neurons. H) Percent colocalization of GFP::SAS-7 with SAS-4::TagRFP: 69.2%, n=79 SAS-7 foci, n=39 NPs, TagRFP::SPD-2: 77.5%, n=80 SAS-7 foci, n=20 NPs. I) Percent colocalization of GFP::SAS-7 with TagRFPT::SAS-1: 51.8%, n=15 SAS-7 foci, n=11 NPs; SAS-4::TagRFP: 67.6%, n=37 SAS-7 foci, n=22 NPs; TagRFP::SPD-2: 63.3%, n=30 SAS-7 foci, n=20 NPs. Since SAS-1 localizes to the transition zone in phasmid cilia ^79^, we only assessed SAS-1 colocalization with SAS-7 foci that were away from the tip of the dendrite to avoid this non-centriolar pool. Scale bars for all images represent 15μm, insets scale bars represent 2μm. All proteins are endogenously tagged with the exceptions of TBG-1, which is a single-copy insertion expressed by the *pie-1* promoter and SAS-4::TagRFP, which is a single-copy insertion driven by the *maph-9* promoter. All images are max projections (unless noted) of images acquired using confocal microscopy of live neurons.

We initially tested CAP-Trap in *C. elegans* embryonic intestinal cells, developing epithelial cells that normally inactivate their centrosomes as MTOCs after the E8-E16 cell division when MTOC function is reassigned to the apical surface ^2,34^. CAP-Trap was successfully targeted to centrioles, as indicated by its colocalization with SAS-7, and led to ectopic activation of MTOC function at the centrosome: centrosomes were associated with microtubules, the PCM protein SPD-5, and the plus end microtubule binding protein EBP-2/EB1, where EBP-2 comets moved away from centrioles (Extended Data Fig. 1a-g, Supplementary Video 1-3). Surprisingly, while only centrioles that clearly localized CAP-Trap were enriched with EBP-2, SPD-5 was expanded at all centrioles in CAP-Trap-expressing intestinal cells, uncoupling SPD-5 expansion from MTOC activation in some cases (Extended Data Fig. 1g). This separation of function is consistent with analysis in *C. elegans* early embryos where mutations in *spd-5* can similarly separate SPD-5 expansion from its role in MTOC function ^31^. In contrast, intestinal centrosomes in embryos expressing CAP-Trap alone, without expression of GFP::SAS-7 (hereafter referred to as ‘CAP’, Extended Data Fig. 2), were not active as MTOCs (Extended Data Fig. 1c), indicating that targeting of CAP to centrioles is essential for effective centrosome activation and in line with previous studies demonstrating that local PLK-1/Polo activity is required to drive centrosome activation in mitosis ^35^. Thus, CAP-Trap can serve as a genetic tool to activate centrosomes in differentiated *C. elegans* cells.

To enable exploratory studies of unpredicted phenotypes caused by CAP-Trap expression, we next sought a cell type that was dispensable for viability and in which roles for microtubules in cell function had been clearly established. We expressed CAP-Trap in *C. elegans* ciliated sensory neurons and focused on the development of the phasmid neurons, two pairs of neurons in the worm tail that are isolated from the majority of other ciliated sensory neurons, enabling single cell analysis of resulting phenotypes (Fig. 1a). The phasmid neurons are born in the embryo ∼270-400 minutes post fertilization (mpf) ^36^. Following this terminal division and during dendrite outgrowth, one of the two centrioles localizes to the dendrite tip and serves as a basal body to template the growth of a primary cilium ^37,38^. After the initiation of ciliogenesis, the centriole is eliminated from the base of each cilium while the centriole in the cell body is eliminated prior to ciliogenesis during initial dendrite extension ^37,39–41^.

We expressed CAP-Trap in the phasmid neurons just after their terminal cell division using the promoter for the *maph-9* gene (*maph-*9p) and assessed the MTOC state of the centrosome in the mature adult neuron. Consistent with published electron microscopy (EM) and high resolution light microscopy data ^19,37,39^, SAS-7 was absent from the cell body and dendrites of control animals due to the programmed elimination of the centriole from this cell type earlier in development (Fig. 1c-e).

We previously found that the base of each cilium becomes an MTOC made from an acentriolar pool of PCM and populates the dendrite with minus end out microtubules^37,39^. In control neurons, we observed the expected enrichment of the PCM component SPD-5, the microtubule plus end binding protein EBP-2/EB1, and the γ-TuRC component TBG-1/γ-tubulin at the tip of the dendrite (Fig. 1c-e). In contrast, CAP-Trap expressing neurons had foci of SAS-7 within the cell body and dendrite that colocalized with SPD-5, TBG-1, and EBP-2, consistent with these foci being active centrosomes (Fig. 1c-e). Indeed, live imaging of EBP-2 showed comets emanating from the SAS-7 positive foci, reflecting microtubule growth from these sites (Fig. 1f, Supplementary Video 4). To confirm that these SAS-7 foci were centrioles, we colocalized more internal components of the centriole (Fig. 1g). In first larval stage (L1) and adult animals expressing CAP-Trap, SAS-7 foci colocalized with SPD-2, SAS-4, and SAS-1 (Fig. 1h, i). Together, these data indicate that CAP-Trap expression leads to the ectopic activation of MTOC function at the centrosome in phasmid neurons.

### Centrosome activation perturbs microtubule polarity and dynein-mediated cargo trafficking

Dendrites in invertebrate neurons have a stereotyped “minus-end-out” microtubule polarity where all microtubules orient with minus ends localized within the dendrite and plus ends pointed towards the cell body (Fig. 2a). In control phasmid neurons, minus-end-out microtubules are nucleated from MTOCs on the membrane at the tip of the dendrite and at the base of cilia ^37,39,42^, as visualized by enriched EBP-2 intensity at the tip of the dendrite and EBP-2 comets moving towards the cell body (Fig. 2c,c’, e, f, Supplementary Video 5). This MTOC activity led to the majority of EBP-2 comets appearing near the tip of the dendrite, as we did not observe much EBP-2 in the cell body or middle of the dendrite (Fig. 2c, c’, e, f). In contrast, CAP-Trap expressing neurons had SAS-7 foci that also localized EBP-2 in the cell body and along the dendritic process. In these cases, EBP-2 comets originated from the cell body or the middle of the dendrite, populating the dendrite with plus end out MTs, some of which entered the cilia region (Fig. 2b, d, d’, Supplementary Video 6). This bidirectional movement of EBP-2 comets demonstrated a clear disruption in the normal dendrite microtubule polarity (Fig. 2d’). Consistently, overall EBP-2 intensity was also increased in CAP-Trap expressing neurons with active centrosomes in the cell body, dendrite process, or at the base of cilia region (Fig. 2e-f,). Thus, centrosome activation disrupts the microtubule polarity in CAP-Trap expressing neurons.

**Figure 2.**
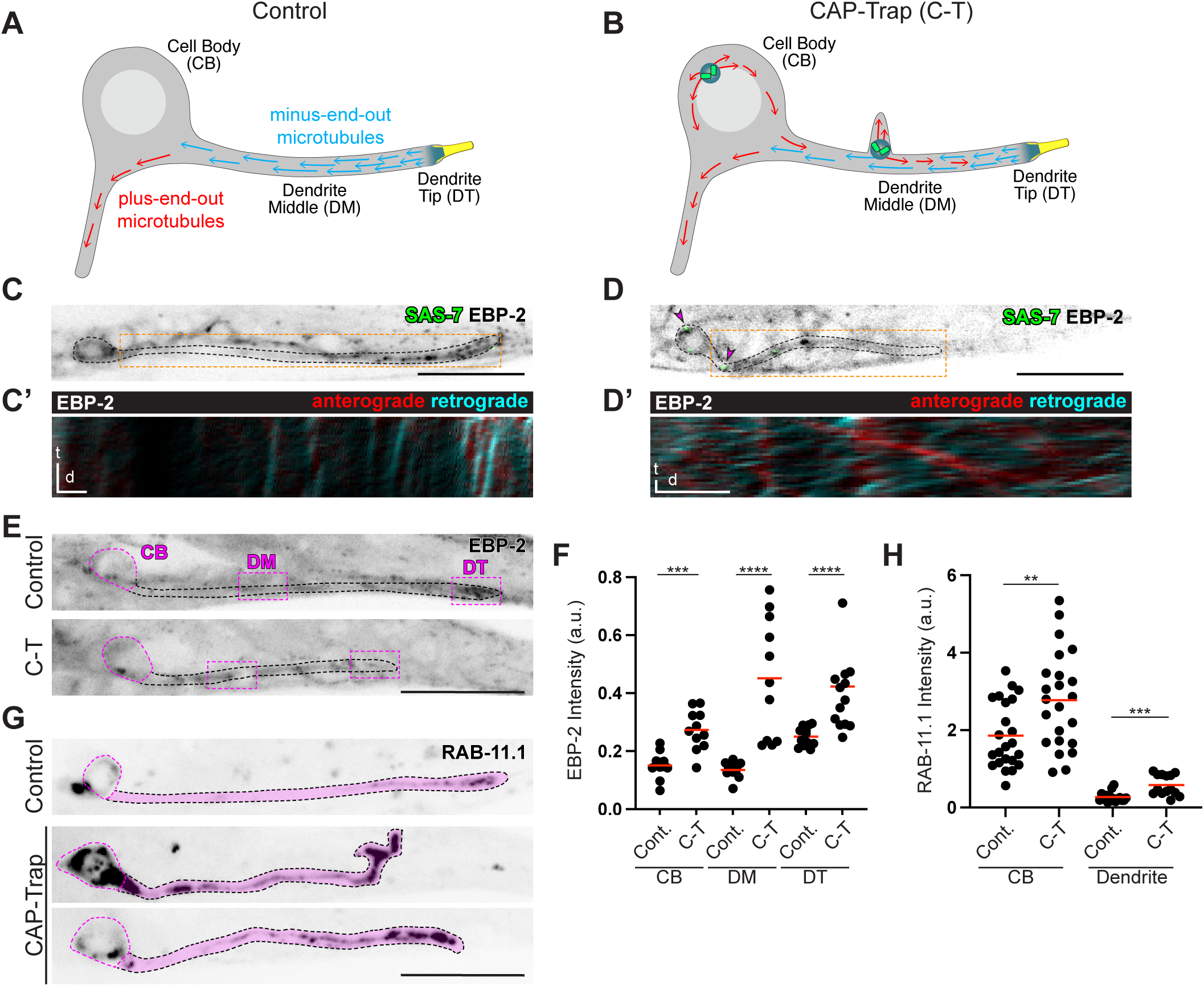
CAP-Trap expression perturbs microtubule polarity and downstream dynein-based cargo trafficking A-B) Schematic of microtubule polarity in controls neurons and in C-T expressing neurons with active centrosomes. Control neurons have minus-end-out microtubules (blue arrows) in the dendrite nucleated from the ncMTOC (teal) at the base of cilia (yellow) at the dendrite tip. Active centrosomes in the cell body and middle of the dendrite nucleate plus end out microtubules (red arrows) into the dendrite in C-T expressing neurons. C-D) Control^BFP-Trap^ and C-T expressing neurons with centrioles (GFP::SAS-7) and MTOC site (EBP-2::mKate-2) labeled. Single-slice images. Arrowheads highlight active centrosomes in C-T expressing neurons. Boxed regions were used to generate the kymographs in C’-D’. C’-D’) Kymographs showing EBP-2 directionality in control and C-T expressing adult neurons. EBP-2 comets were filtered based on directionality and color-coded based on anterograde or retrograde movement (see Methods). Kymographs were generated using single-slice time lapse movies of EBP-2 dynamics ( d=2μm , t=3.7 sec for control, t=10 sec for C-T). E) Adult Control^No C-T^ and C-T expressing neurons with EBP-2::mKate-2 labeled. Regions outlined in magenta represent the cellular regions used for EBP-2 intensity quantifications. F) Quantification of overall EBP-2 intensity in Control^No^ ^C-T^ and C-T expressing neurons (Control CB (‘Cell Body’): n=12 NPs, C-T CB: n=11 NPs, Control DM (‘Dendrite Middle’): n=13 NPs, C-T DM: n=11 NPs, Control DT (‘Dendrite Tip’): n=14 NPs, C-T DT: n=12 NPs). G) Distribution of neuronal-specific mScarlet::RAB-11.1 in adult Control^No C-T^ and C-T expressing neurons. The magenta regions represent the cellular regions used to quantify RAB-11.1 intensity. H) Quantification of total RAB-11.1 intensity in the cell body and dendrite in Control^No C-T^ and C-T expressing neurons (Control CB: n=23 NPs, C-T CB: n=23 NPs, Control Dendrite: n=16 NPs, C-T Dendrite: n=14 NPs). Scale bars for all images represent 15μm, with the exception of the kymographs in C’-D’. Quantification: red line: mean, Mann-Whitney U test. p values: **=p ≤ 0.01, ***=p ≤ 0.001, ****=p ≤ 0.0001. All proteins are endogenously tagged. All images are max projections of images acquired using confocal microscopy of live neurons.

Stereotyped microtubule polarity is essential in neurons for the polarized trafficking of cargo to and from axons and dendrites. Specifically, minus-end-out microtubule polarity in dendrites allows dynein-mediated trafficking of cargo to the tip of the dendrite. In control neurons, the dynein cargo RAB-11.1 shows an enrichment in the distal parts of the phasmid dendrite. Consistent with changes in dendrite microtubule polarity following CAP-Trap expression, RAB-11.1 was highly enriched in large aggregates throughout the length of the dendrite and in the cell body (Fig. 2g-h). Thus, centrosome activation through CAP-Trap expression generates new MTOC sites that perturb the stereotyped microtubule polarity and trafficking in neurons.

### Centrosome activation perturbs dendrite morphology

Having demonstrated the effectiveness of CAP-Trap as a tool to activate centrosomal MTOC capacity, we sought to determine whether the resulting change in microtubule organization impacted cell function beyond cargo trafficking. Since CAP-Trap overexpresses active PLK-1 throughout the cell, which may itself influence cellular processes independent of centrosome activation, we designed a series of controls to confirm that observed phenotypes were due to the role of CAP-Trap at the centrosome rather than elsewhere in the cell. In particular, we used a CAP control, in which the same experimental CAP transgene was expressed in a background in which SAS-7 was untagged, leaving CAP localized throughout the cytoplasm rather than targeted to the centrioles (Extended Data Fig. 2). We expected some low level of centrosomal MTOC activation in these CAP expressing control neurons, but to a significantly lesser extent than in those expressing CAP-Trap. In addition to the CAP control, we also compared CAP-Trap phenotypes to a control strain in which CAP-Trap was not expressed (“no CAP-Trap”) or in which BFP fused to the GFP nanobody was expressed from the same promoter (“BFP-Trap”, Extended Data Fig. 2).

Using this experimental framework, we assessed neuronal development and found severe dendrite morphology defects in CAP-Trap expressing neurons (Fig. 3a). Control phasmid neurons displayed the stereotypical morphology with an axon extending ventrally from the cell body, and a posteriorly extended dendrite. In contrast, CAP-Trap expressing phasmid neurons displayed severe dendrite morphology defects in which dendrites were short, contained ectopic protrusions, and were occasionally oriented towards the anterior of the worm (Fig. 3a). Dendrite length in adult animals was much shorter following CAP-Trap expression than in control phasmid neurons (Fig. 3b). This dendrite length defect was present throughout development, even in L1 larvae (Fig. 3e-f). Surprisingly, in 75.0% of phasmid dendrites in CAP-Trap and 38.7% in CAP expressing L1s, the dendrite appeared fragmented with a residual membrane associated structure that also contained a SAS-7 focus (Fig. 3e), likely representing the detachment of the dendrite tip containing a centriole.

**Figure 3.**
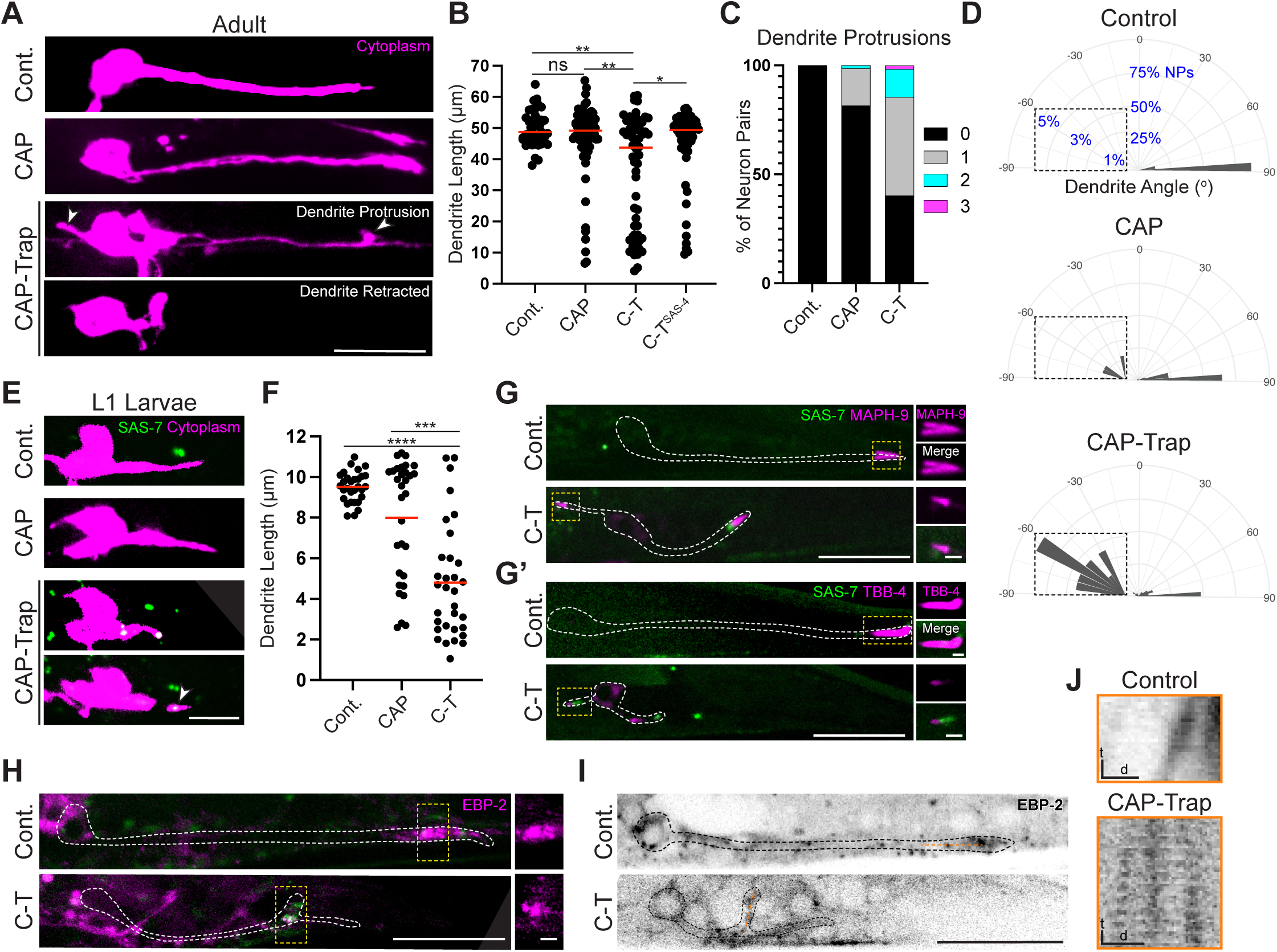
CAP-Trap expression perturbs dendrite morphology A) Morphology of adult Control^BFP-Trap^, CAP, and C-T expressing neurons. Control and CAP expressing neurons have posteriorly extended dendrites. C-T neurons have backward or non-extended dendrites and dendrites with ectopic protrusions (arrowhead). Images were scaled to highlight dendrite morphology. B) Quantification of dendrite protrusions in adult neurons (Control^BFP-Trap^: n=40 NPs, CAP: n=76 NPs, C-T: n=60 NPs). C) Quantification of dendrite length in adult neurons of indicated genotypes (Control^BFP-Trap^: n=40 NPs, CAP: n=76 NPs, C-T: n=60 NPs, red line: median, Kruskal-Wallis test followed by Dunn’s multiple comparisons test). C-T^SAS-4^ neurons express SAS-4::GFP instead of GFP:SAS-7. (C-T^SAS-4^ n=60 NPS, red line: median, Mann-Whitney U test). D) Dendrite angles for adult neurons of indicated genotypes. Insets highlight backward protrusions (Control^BFP-Trap^: n=39 angles, CAP: n=82 angles, C-T: n=79 angles, Kruskal-Wallis test followed by Dunn’s multiple comparisons). E) Dendrite morphology of Control^BFP-Trap^, CAP, and C-T expressing L1 larvae. Scale bar: 5μm. Centrioles (GFP::SAS-7) are retained in C-T animals, some of which are retained in a broken-off piece of dendrite (arrowhead) (CAP: 38.7 % of NPs with broken-off dendrite, n=31 NPs; C-T: 75.0% of NPs with broken-off dendrite, 91.3% of dendrite pieces retain SAS-7, n=32 NPs). F) Quantification of dendrite length in L1 neurons (Control^BFP-Trap^: n=30 NPs, CAP: n=31 NPs, C-T: n=32 NPs, red line: median, Kruskal-Wallis test followed by Dunn’s multiple comparisons test). G-H) Control^No C-T^ and C-T expressing adult neurons with centriolar GFP::SAS-7 and ciliary doublet protein TagRFP::MAPH-9 (G), ciliary singlet protein TBB-4::RFP (G’), or EBP-2::mKate2 (H, I). Magnified insets of boxed regions shown at right. Control animals have dendrites extended posterior to the cell body with cilia at the dendrite tip (n=21/21 NPs), while C-T expressing neurons have backward ciliated dendrites extending to the anterior (n=19/26 NPs). H) In control neurons, EBP-2 is enriched at the tip of the dendrite (n=56/56 NPs). In C-T animals, SAS-7 foci that recruit EBP-2 are enriched at dendrite protrusions (88.0% of protrusions with SAS-7, 85.7% of SAS-7 foci with EBP-2; n=25 NPs). I) Orange line represents the region used to generate the kymographs in J. J) Kymograph of EBP-2 dynamics in adult Control^No C-T^ neurons and in dendritic protrusions in C-T expressing neurons. Kymographs were generated using single-slice movies of EBP-2 dynamics. Scale bars: t=2sec, d=2μm. p values: ns=p>0.05, *=p ≤ 0.05, **=p ≤ 0.01, ***=p ≤ 0.001, ****=p ≤ 0.0001. All proteins are endogenously tagged. All images are max projections of images acquired using confocal microscopy of live neurons. Images were scaled to better show dendrite morphology. Unless otherwise indicated, scale bar: 15μm, Inset scale bar: 2μm.

In addition to dendrite length defects, CAP-Trap expressing neurons also contained ectopic protrusions in adult animals (Fig. 3c). Intriguingly, SAS-7 and EBP-2 were consistently enriched at the base of each protrusion (Fig. 3h, Supplementary Video 7), and live imaging showed EBP-2 comets moving into the protrusions, consistent with centrosomal MTOC activity generating and/or stabilizing these protrusions (Fig. 3i-j). We also found that dendrite angles were highly variable in CAP-Trap expressing phasmids compared to CAP expressing or control phasmids. While control dendrite angles were generally 90°, representing the normal posterior extension of the dendrite in relation to the cell body, CAP-Trap expressing neurons had severe dendrite pathfinding defects with dendrite angles that ranged from 34° to -89°, and some dendrites pointing anteriorly from the cell body (Fig. 3d). To determine if these anterior processes were dendrites, we assessed the localization of the cilia markers MAPH-9 and TBB-4, which are normally localized at the posterior end of the dendrite in control neurons (Fig. 3g-g’). In CAP-Trap expressing phasmids, 73.1% of processes projecting anterior to the cell body localized cilia components, confirming that some of these structures were indeed backwards dendrites (Fig. 3g-g’).

To further confirm that dendrite morphology defects were due to centrosomal targeting of PLK-1, rather than PLK-1 overexpression *per se*, we targeted CAP to a more internal position on the centriole using GFP::SAS-4 ^43^, reasoning that this internal targeting should be less efficient at reactivating the MTOC function of the centrosome. Indeed, targeting CAP to SAS-4 reduced dendrite morphology defects compared to targeting with SAS-7 (Fig. 3b). This ability to tune the percentage of morphology defects further supports our claim that the morphology defects were due to the ability of CAP-Trap to activate MTOC function at the centrosome, rather than effects of PLK-1 elsewhere in the cell.

Finally, we wanted to determine if the dendrite morphology phenotypes observed following CAP-Trap expression were specifically due to the ectopic MTOC activity of the centrosome. We hypothesized that degrading PCM proteins in CAP-Trap expressing neurons should suppress such phenotypes, as the ability of the centrosome to build microtubules would be eliminated. To test this hypothesis, we endogenously tagged the main centrosomal MTOC component SPD-5 with the ZF degron and tissue-specifically expressed the E3 ligase adapter ZIF-1 in post-mitotic ciliated sensory neurons (csn) using the promoter for the gene *osm-6* (SPD-5^csn(-)^) ^44–46^. SPD-5 degradation suppressed the dendrite length, protrusion, and angle phenotypes in CAP-Trap expressing phasmids (Fig. 4a-e), indicating that these dendrite morphology defects were due to centrosomal MTOC activity. To completely eliminate PCM, we removed the three main PCM proteins in *C. elegans,* SPD-2, SPD-5, and PCMD-1 (‘PCM^csn(-)^’), in CAP-Trap expressing phasmid neurons using the same ZF/ZIF-1 degradation strategy. Strikingly, we found that the dendrite length phenotype was further suppressed in PCM^csn(-)^ phasmids compared to SPD-5^csn(-)^ alone (Fig. 4f-g). The ability of PCM degradation to suppress CAP-Trap phenotypes demonstrates that ectopic centrosome activation causes severe neuronal morphology defects.

**Figure 4.**
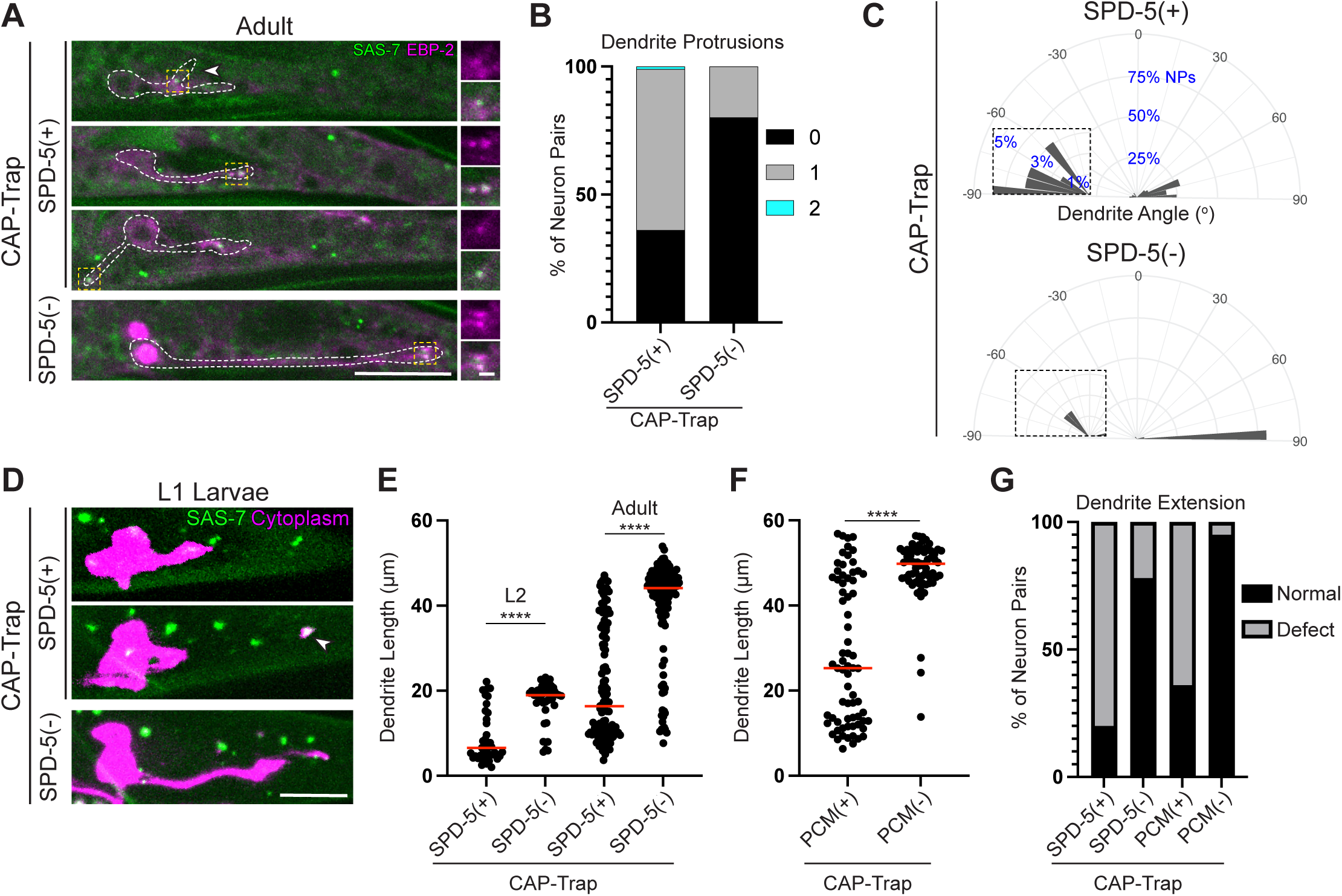
Centrosomal MTOC activity perturbs dendrite morphology A) C-T expressing adult neurons with (SPD-5^(-)^) or without (SPD-5^(+)^) SPD-5 degraded only in ciliated sensory neurons (SPD-5^csn(-)^). Centrioles (GFP::SAS-7) and microtubule plus ends (EBP-2::mKate2) are labeled. SPD-5^(+)^ neurons have backwards dendrites, dendrites that do not extend, and dendrite protrusions (arrowhead). Magnified insets of boxed regions shown at right. Scale bar: 15μm, inset scale bar: 2μm. B) Quantification of dendrite protrusions in adult neurons (SPD-5^(+)^: n=79 NPs, SPD-5^(-)^: n=69 NPs). C) Dendrite angles of adult neurons. Insets highlight backward dendrites (SPD-5^(+)^: n=65 angles, SPD-5^(-)^: n=65 angles), Mann-Whitney U test:****. D) Dendrite morphology in L2 neurons. SPD-5^(+)^ neurons have non-extended dendrites with broken-off pieces that contain GFP::SAS-7 (arrowhead). Images were scaled to better show dendrite morphology. Scale bar: 5μm. E) Quantification of dendrite length in L2 and adult neurons (L2: SPD-5^(+)^: n=41 NPs; SPD-5^(-)^: n=46 NPs, adult: SPD-5^(+)^: n=122 NPs; SPD-5^(-)^: n=121 NPs), red line: median, Mann-Whitney U test). Scale bar: 5μm. F) Quantification of dendrite length in C-T expressing neurons with (PCM^(-)^) and without (PCM^(+)^) PCM components SPD-5, SPD-2, and PCMD-1 degraded only in ciliated sensory neurons (PCM^csn(-)^). (PCM^(+)^ n=70 NPs, PCM^(-)^ n=65 NPs). Red line: median. Mann-Whitney U test. G) Quantification of dendrite length in adult neurons (SPD-5^(+)^: n=122 NPs; SPD-5^(-)^: n=121 NPs, PCM^(+)^ n=70 NPs; PCM^(-)^ n=65 NPs). p values:****=p ≤ 0.0001. All proteins are endogenously tagged. All images are max projections of images acquired using confocal microscopy of live neurons.

### Ectopic PCM perturbs ciliogenesis

A main cellular function for the centriole is to act as a basal body to template the assembly of a cilium. In differentiating cells where centrosomes are inactivated, basal bodies associate with little or no PCM ^47,48^, and thus, their ectopic activation as an MTOC is predicted to severely perturb ciliogenesis. This should especially be true for *C. elegans* cilia, where centrioles are normally eliminated from the ciliary base following the onset of ciliogenesis and cilia instead recruit the PCM protein SPD-5 inside the base of the axoneme to produce a new acentriolar MTOC ^37,39^. Thus, we hypothesized that CAP-Trap expression should surround the normally naked ciliary base with PCM and therefore potentially obstruct aspects of ciliogenesis.

As an initial assessment of potential cilia defects, we used a “dyefill assay” that tests the ability of externally exposed cilia to take up a lipophilic dye from the environment. This assay is a crude measure of cilia defects as cilia that are structurally perturbed are often defective in taking up dye ^38,49^. We found that almost all CAP and CAP-Trap expressing animals were dyefill defective, regardless of dendrite length (Fig. 5a-b). To more specifically probe overall cilia structure and function in CAP-Trap expressing neurons, we localized the microtubule doublet protein MAPH-9/MAP9 and TBB-4/β-tubulin that is enriched in the singlet regions of the axoneme. Surprisingly, we found that most CAP-Trap expressing neurons were ciliated, regardless of dendrite morphology, although both MAPH-9 and TBB-4 were both reduced in length and intensity compared to in control (Fig. 5d-i). While MAPH-9 was detected in 94.3% of neurons, TBB-4 was not detected in 62.5% of neurons, indicating that the singlet region was entirely missing in these CAP-Trap expressing neurons. In rare cases where neurons had no detectable cilia, we found a focus of MAPH-9 in the cell body, consistent with the presence of a centriole that never converted to a basal body (Fig. 5j). Thus, CAP-Trap expressing neurons have cilia that are structurally and functionally defective.

**Figure 5.**
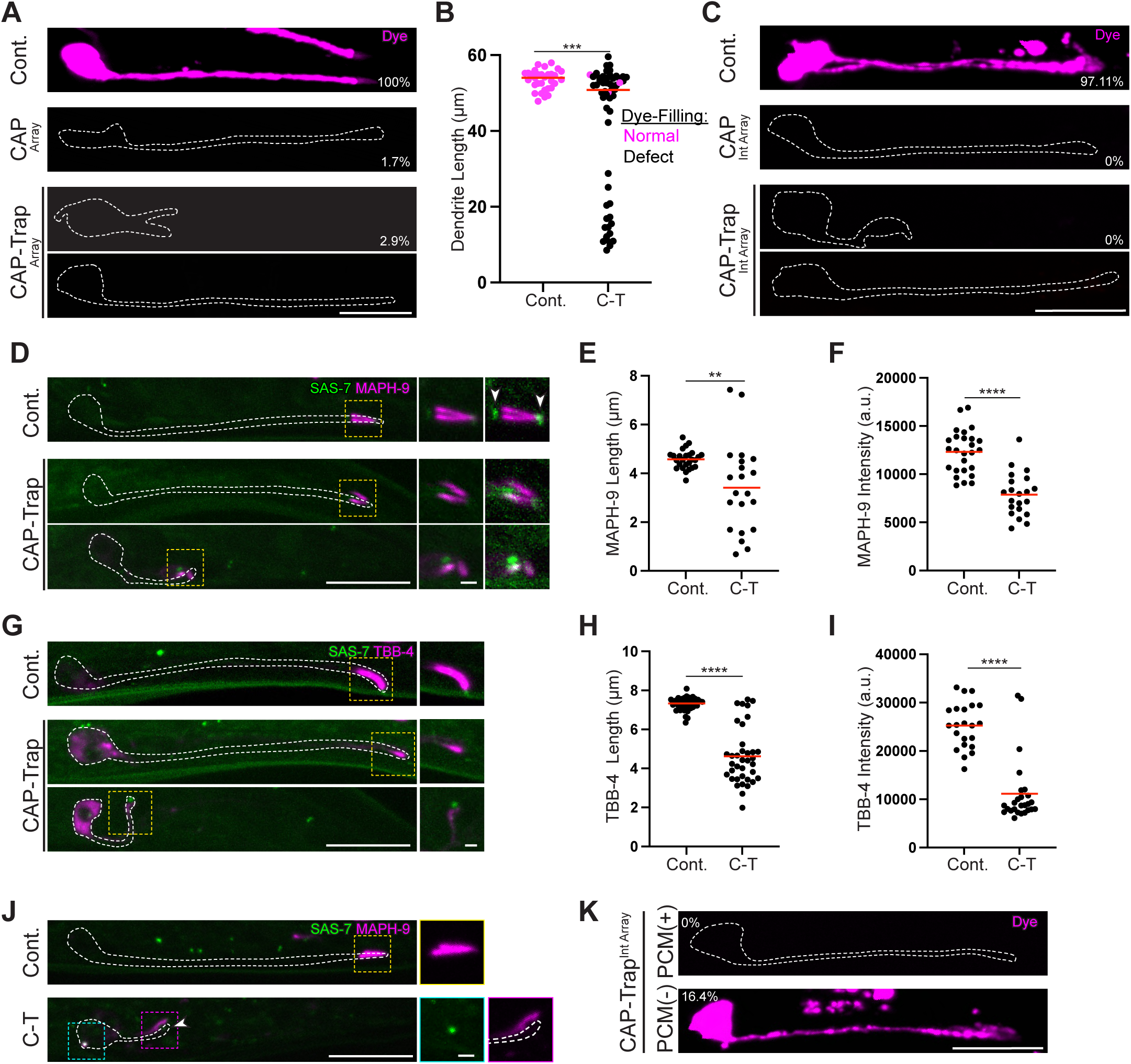
CAP-Trap expression perturbs cilia structure and function A) Dyefill experiment in Control^No CT^, CAP^Array^, and C-T^Array^ expressing animals. Percentages correspond to the number of neurons that dye-fill. (Control^No C-T^: n=60 neurons, CAP^Array^: n=59 neurons, C-T^Array^: n=103 neurons). B) Quantification of dendrite length in Control^No CT^ and C-T^Array^. The majority of C-T expressing neurons did not dyefill regardless of whether the dendrite length was defective (Control: n=32 NPs, C-T: n=56 NPs). C) Dyefill experiment in Control^BFP-Trap^ , CAP^Int Array^, C-T^Int Array^ expressing neurons (Control^BFP-Trap^: n=72 neurons, CAP^Int Array^: n=88 neurons, C-T^Int Array^: n=88 neurons). Percentages correspond to the number of neurons that dye-filled for each genotype. D) Control^No C-T^ and C-T expressing neurons with centrioles (GFP::SAS-7) and ciliary microtubule doublet marker (TagRFP::MAPH-9) labeled. Note that SAS-7 often localized around the periciliary membrane compartment proximal to the tip of the dendrite, likely within the glia that surround each neuron. LUTs were re-scaled in the second set of insets and arrowheads added to highlight this non-centriolar localization of GFP::SAS-7 in control animals. 3/53 neurons lack MAPH-9. E-F) Quantification of MAPH-9 length and intensity in Control^No C-T^ and C-T expressing neurons (Control: 22 neurons, C-T: 26 neurons). G) Control^No C-T^ and C-T expressing neurons with centrioles (GFP::SAS-7) and ciliary microtubule singlet marker (TBB-4::Tag::RFP) labeled. 21/56 C-T expressing neurons lack TBB-4. H-I) Quantification of TBB-4 length and intensity. TBB-4 length: Control: 41 neurons, C-T: 37 neurons. TBB-4 intensity: Control: 22 neurons, C-T: 26 neurons. J) Control and C-T expressing neurons with centrioles (GFP::SAS-7) and MAPH-9 labeled. In cases where MAPH-9 is localized to the centriole in the cell body, no enrichment of MAPH-9 is detected at the tip of the dendrite where the cilium would normally localize (n=4 NPs). K) Dyefill experiment in C-T expressing neurons with (PCM^(-)^) or without PCM degraded (PCM^(+)^) only in ciliated sensory neurons (PCM^csn(-)^). Percentages correspond to the number of neurons that dye-filled. (PCM^(+)^: n=116 neurons, PCM^(-)^: n=122 neurons). Scale bars all represent 15μm with insets representing 2μm. Quantification: Mann-Whitney U test. p values: **=p ≤ 0.01, ***=p ≤ 0.001, ****=p ≤ 0.0001. All proteins are endogenously tagged. All images are max projections of images acquired using confocal microscopy of live neurons. Magnified insets of boxed regions shown at the right.

To determine whether cilia defects following CAP-Trap expression were due to the ectopic reactivation of the centrosome, we tested whether the removal of PCM proteins from the phasmid neurons (PCM^csn(-)^) could suppress the ciliogenesis phenotypes seen following CAP-Trap expression. In a strain where CAP-Trap is stably integrated into the genome, 0% of CAP or CAP-Trap expressing neurons dye fill (Fig. 5c). Surprisingly, we found that 16.4% of PCM^csn(-)^, CAP-Trap expressing neurons dye-filled, indicating a suppression of the ciliogenesis defects (Fig. 5k). Thus, ectopic PCM caused by CAP-Trap expression contributes to defects in ciliogenesis, although ectopic PLK-1 overexpression may also contribute to this phenotype.

### Dendrite length phenotypes are correlated with defects in the cilia transition zone

In the phasmid neurons, dendrites elongate through retrograde extension where the cilium in each dendrite is stably adhered to glial cells at the posterior while the cell body migrates anteriorly, thus extending the dendrite ^50,51^. We therefore reasoned that the dendrite extension defects we observed in CAP-Trap expressing neurons could be due to cilia defects, however we needed to rule out that defects in the cilia environment did not instead lead to perturbed dendrite extension. The phasmid neurons are encased by a sheath cell and each cilium projects into a pore created by two glial socket cells that opens to the environment (Fig. 6a). To assess the phasmid pore, we again turned to the dyefill assay, reasoning that only cilia surrounded by an intact pore could take up dye. In mosaic animals in which CAP-Trap was expressed only in one of the two neurons in the pair, the neuron without CAP-Trap expression was able to dye fill, indicating that the shared pore structure was likely unperturbed (Fig. 6b). To further assess the pore, we assayed cell-cell adhesions within the pore which is formed from two glial socket cells that each wrap and form an autojunction and adhere to one another (Fig. 6a). High resolution imaging showed that this arrangement results in a line of the homophilic adhesion protein HMR-1/E-cadherin posterior to the base of cilia that ends in two concentric rings at the posterior opening of the pore (Fig. 6a’). In control animals, this localization appeared as two enrichments of HMR-1 at these sites of adhesion—one at the line of adhesion and one at the concentric rings (Fig. 6c). In mosaic animals in which CAP-Trap was only expressed in one of the four phasmid neurons, HMR-1 in the CAP-Trap expressing neuron appeared unperturbed compared to in control (Fig. 6c). Similarly, in both retracted and extended dendrites in CAP-Trap expressing animals, adhesion was present and at similar intensity as in controls, indicating that the pore structure was intact (Fig. 6e-f). In contrast, we found that HMR-1 length was reduced in CAP-Trap expressing neurons that did not extend compared to control and extended CAP-Trap neurons (Fig. 6g), perhaps indicating a defect in the boundary between the socket and sheath cells. Thus, the pore structure is likely intact in CAP-Trap expressing neurons, pointing to a specific defect in cilia structure that perturbs dendrite extension.

**Figure 6.**
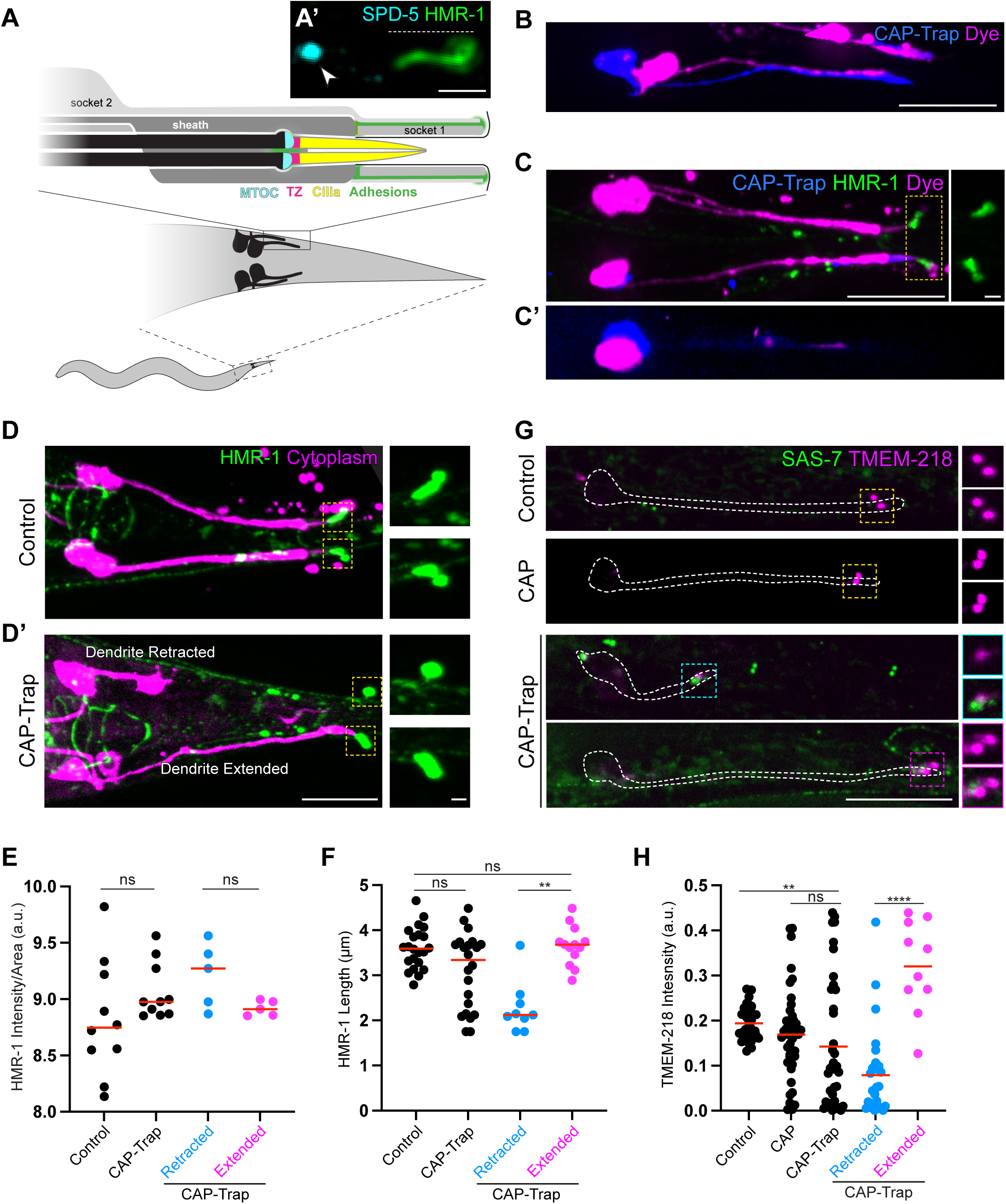
Dendrite extension defects are correlated with defects in the cilia transition zone A) Schematic of the two pairs of phasmid neurons in the *C*. *elegans* tail. Interactions between the phasmid neuron dendrite with the surrounding glial cells (one sheath cell and two socket cells) with sites of adhesion highlighted (purple). A’) High resolution microscopy image of the base of cilia (TagRFP::SPD-5) and adherens junction (HMR-1/E-cadherin::GFP. Arrowhead indicates the base of cilia MTOC and the dotted line indicates HMR-1 adhesion in the phasmid pore region of an adult Control^No C-T^ neuron. The HMR-1 and SPD-5 channels were individually re-sliced and combined in Fiij to highlight the HMR-1 ring structure at the pore opening. Scale bar: 2μm. B) Pair of adult phasmid neurons expressing C-T mosaically with dye (magenta) and C-T (blue) labeled. The neuron without C-T expression dye-fills. C) Two pairs of adult phasmid neurons expressing C-T mosaically with adhesion (HMR-1), dye, and C-T labeled. The adhesion site around the pore in Control (top pair and one neuron in bottom pair) and C-T expressing neurons (one neuron in bottom pair) appear similar. Individual channels were re-sliced and combined to better highlight the presence of the two mosaic cells in the bottom neuron pair. C’) Orthogonal view highlighting that only one of the two neurons in the bottom neuron pair expresses C-T and does not dye fill. D) Control^No^ ^C-T^ and C-T expressing neurons with the cytoplasm and HMR-1 labeled. D’) C-T expressing neurons with retracted (top) and extended dendrites (bottom). E-F) Quantification of HMR-1 intensity and length. HMR-1 intensity in Control^No CT^ (n=10 NPs) and C-T expressing neurons (n=10 NPs) with retracted (n=5 NPs) or extended (n=5 NPs) dendrites. Red line: median. Quantification of HMR-1 length in Control^No C-T^ (n=22 NPs) and C-T expressing neurons (n=22 NPs) with retracted (n=9 NPs) or extended (n=13 NPs) dendrites. Red line: median. G) Localization of GFP::SAS-7 and the transition zone component TMEM-218 in Control^No C-T^, CAP, and C-T expressing neurons with retracted (top) and extended (Bottom) dendrites. H) Quantification of TMEM-218 intensity (Control: n=34 neurons, CAP: n=42 neurons, C-T: n=38 neurons, C-T retracted: n=28 neurons, C-T extended: n=10 neurons, red line: median). Quantification: Kruskal-Wallis test followed by Dunn’s multiple comparisons test. p values: ns=p>0.05, **=p ≤ 0.01,****=p ≤ 0.0001. All proteins are endogenously tagged. All images are max projections of images acquired using confocal microscopy of live neurons. Magnified insets of boxed regions shown at right. Scale bars: 15μm, inset scale bars: 2μm.

A specialized region of the cilium known as the transition zone (TZ) is thought to mediate adhesion between cilia and the surrounding glial sheath cell since TZ mutants have retracted dendrites due to a failure to attach to the sheath during retrograde extension ^51–54^. Consistent with a role for the TZ in dendrite extension, we found that CAP-Trap expressing phasmids had severe defects in the localization of the TZ component TMEM-218 compared to in control and CAP expressing neurons (Fig. 6g). TMEM-218 intensity was consistently lower in retracted dendrites (Fig. 6h). These data are consistent with a model where ectopic MTOC function at the centrosome perturbs the TZ, thereby leading to defects in ciliary adhesion and dendrite extension.

### Ectopic PCM protects centrioles from elimination

Centrioles are eliminated through unknown mechanisms in a variety of differentiated cell types such as in striated muscle cells and oocytes across organisms ^20^. Centriole elimination in *C. elegans* is stereotyped, with their removal in ciliated neurons shortly after the beginning of ciliogenesis during embryonic development ^19,41^. We were therefore surprised to consistently observe centriolar foci that colocalized SAS-7 and other centriolar components (Fig. 1h-i) in the ciliated sensory neurons of adult CAP-Trap expressing animals (Fig. 7a-b). These foci were mostly found in the cell body and at the tip of the dendrite, where the centriole would have been prior to normal elimination in control animals (Fig. 7c). Furthermore, we observed a higher percentage of colocalization between SAS-7 and other centriolar markers in CAP-Trap expressing phasmids in L1 larvae compared to adults, indicating that CAP-Trap expression likely delays centriole elimination (Fig. 1h-i). This apparent protection from centriole elimination was not restricted to the phasmids as we also observed retention of SAS-7 foci in the head amphid neurons in both embryos and larvae (Extended Data Fig. 3a-c). Thus, CAP-Trap expression prevents physiological centriole elimination in sensory neurons.

**Figure 7.**
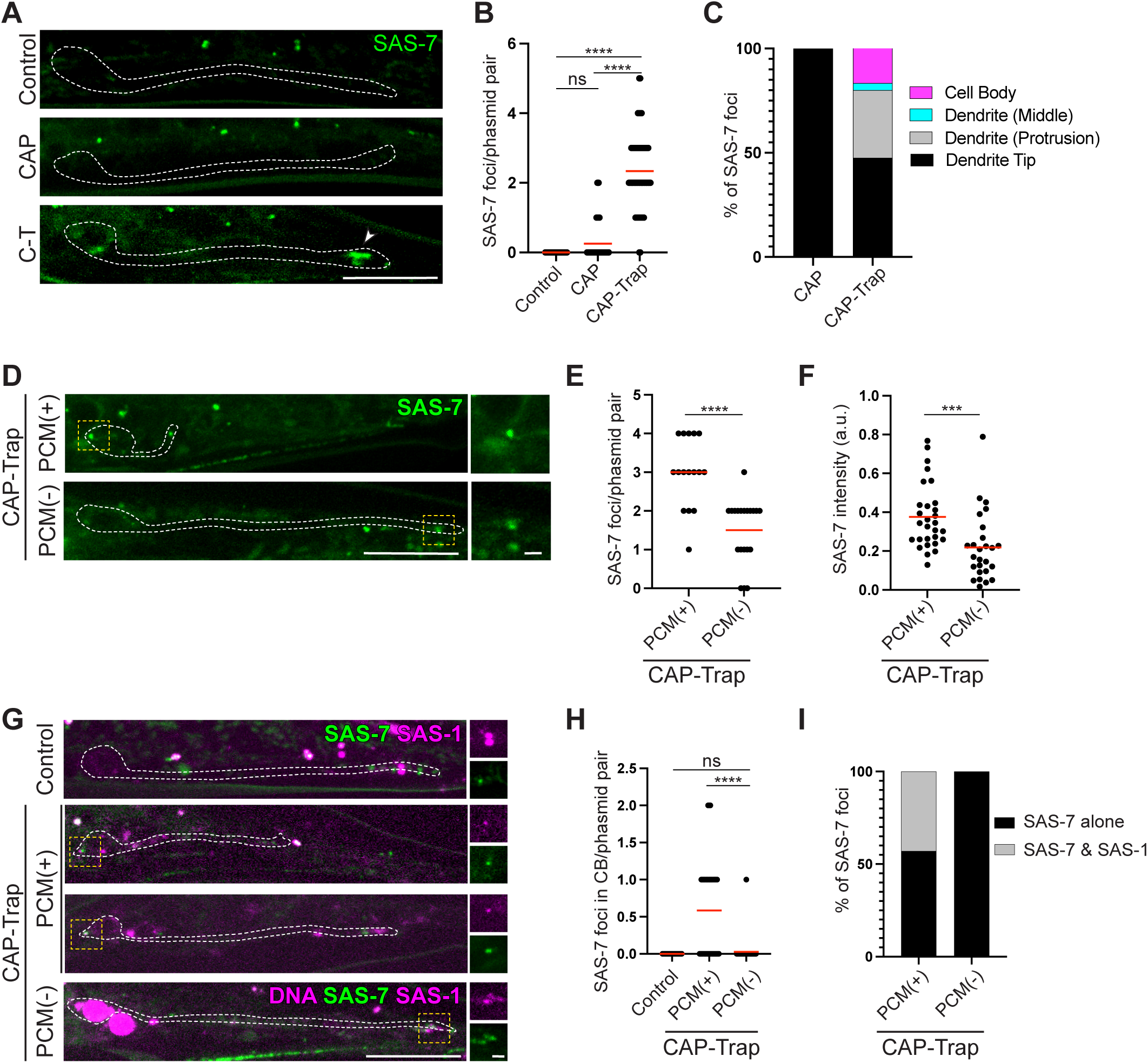
Ectopic PCM protects centrioles from elimination A) Control^No C-T^ and C-T expressing adult neurons express GFP::SAS-7 while CAP neurons express mStayGold::SAS-7. B) Quantification of SAS-7 foci per neuron pair (Control: 39 NPs, CAP: 63 NPs, C-T: 62 NPs; Red line: mean. Kruskal-Wallis test followed by Dunn’s multiple comparisons test). C) Distribution of SAS-7 foci within the neuron (CAP: 63 NPs, C-T: 62 NPs). D). Localization of GFP::SAS-7 in C-T expressing neurons with (PCM^(-)^) and without (PCM^(+)^) PCM components SPD-5, SPD-2, and PCMD-1 degraded only in ciliated sensory neurons (PCM^csn(-)^). E-F) Quantification of the number (E) and intensity (F) of SAS-7 foci in PCM^(+)^ (n=16 NPs) and PCM^(-)^ (n=20 NPs) C-T expressing adult neurons. Red line: mean. Mann-Whitney U test. G). Localization of SAS-1 in Control^No CT^ and C-T expressing neurons in PCM^(+)^ and PCM^(-)^ conditions. PCM is degraded only in ciliated sensory neurons (PCM^csn(-)^). In some cases, SAS-7 and SAS-1 do not perfectly colocalize, likely reflecting SAS-1 localization to the transition zone of cilia (arrowhead) ^79^. H) Quantification of the number of SAS-7 foci in the cell body in Control^No CT^ (n=32 NPs), PCM^(+)^ (n=36 NPs) and PCM^(-)^ (n=36 NPs) C-T expressing neurons. Red line: mean. Kruskal-Wallis test followed by Dunn’s multiple comparisons test. I) Colocalization of SAS-1 with SAS-7 foci in the cell body of PCM^(+)^ (n=21 SAS-7 foci, n=36 NPs) and PCM^(-)^ (n=1 SAS-7 foci, n=36 NPs) C-T expressing neurons. p values: ns=p>0.05, ***=p ≤ 0.001, ****=p ≤ 0.0001. All proteins are endogenously tagged. All images are max projections of images acquired using confocal microscopy of live neurons. Magnified insets of boxed regions shown at the right. Scale bar: 15μm; Inset scale bars: 2μm.

A previous study using *Drosophila* cultured cells suggested that excess PCM surrounding centrioles could inhibit centriole elimination ^21^. To determine whether PCM protects centrioles from elimination *in vivo*, we tested whether removal of PCM in our PCM^csn(-)^ strain would suppress the centriole elimination defect we observed in CAP-Trap expressing neurons. Indeed, we found that the number and intensity of SAS-7 foci were reduced in PCM^csn(-)^, CAP-Trap-expressing neurons compared to neurons without PCM degraded (PCM^csn(+)^) (Fig. 7d-f). We then assessed the colocalization between SAS-7 and SAS-1 only in SAS-7 foci retained in the cell body, away from the cilia, and found that while PCM^csn(+).^ animals retained SAS-7 foci that colocalized with SAS-1, PCM^csn(-)^ animals did not (Fig. 7g-i). Thus, upon CAP-Trap expression, ectopic PCM at active centrosomes protects centrioles from normal elimination during phasmid development, revealing a mechanism that can prevent centriole elimination in a fully differentiated cell *in vivo*.

## Discussion

Cell differentiation requires the rearrangement of intracellular structures to enable specific cell functions, a process that is intimately linked to the dramatic remodeling of the microtubule cytoskeleton. Here, using CAP-Trap to ectopically activate the MTOC function of centrosomes in *C. elegans* phasmid neurons, we challenged the normal non-centrosomal organization of microtubules in this cell type, not only revealing roles for centrosome inactivation in neuronal development, but also highlighting novel microtubule functions in neurons.

CAP-Trap expression caused severe dendrite morphology defects, raising the question of how an active centrosome can perturb neuronal development. For example, we observed dendritic protrusions with active centrosomes at their base upon CAP-Trap expression that were suppressed following SPD-5 degradation, indicating some role for microtubules in protrusion formation. These protrusions appear similar to dendritic varicosities in *Drosophila* da neurons or basal varicosities in mammalian radial glial cells in which microtubules also originate from ncMTOCs ^55,56^. Although microtubules have not been proposed to generate protrusions in these other cell types, our work might reveal a role for microtubules in membrane deformation and/or maintenance and would be consistent with the ability of microtubules to deform membranes *in vitro* ^57^. Although future work will be needed to explore a role for actin, such dendritic protrusions could also form using a similar mechanism as in dendritic growth cones, which rely on the interplay between actin and microtubules ^8,58,59^.

As perhaps the most extreme example of this phenomenon, we occasionally observed the presence of backward-facing dendrites following CAP-Trap expression, suggesting that dendrites were initiated and/or stabilized in the wrong direction. This type of pathfinding defect could be indicative of a defect in the initial establishment of cell polarity, a process that has long been reported to require microtubules in other types of neurons ^60,61^. For example, although several neurites are formed initially in cultured neurons, only certain neurites are selected as the future axon and dendrites, mostly through stabilization of the growth cone through actin and microtubule bundles ^61,63^. Centrosomes have also been directly implicated in this stabilization process, as centrosomes have been found associated with and determine the site of axon outgrowth^63^. The ectopic activation of MTOC activity at the centrosomes in CAP-Trap expressing neurons could similarly misspecify the site of neurite outgrowth in phasmid neurons and/or stabilize neurites protruding in the wrong direction, leading to backwards dendrites. This dendrite mispositioning could also occur through a more passive process, in which dendritic tips that fail to adhere properly at the dendrite pore make inappropriate adhesions that would pull the dendrite backwards during retrograde extension.

The most prevalent phenotype we observed in CAP-Trap expressing neurons was a defect in dendrite extension which correlated with a defective TZ. A systematic study in *Drosophila* ciliated cells showed that the composition of PCM around the basal bodies is specific and varies across cell types ^64^. CAP-Trap likely mimics mitotic PCM and thus would perturb any specialized interphase PCM that might exist in this cell type that could be necessary to convert the centriole to a proper basal body to build a TZ. Excess PCM and/or microtubules around the centriole could also directly impede the recruitment of TZ components. In most systems, TZ formation requires CEP290 and the recruitment of proteins of the MKS and NPHP modules ^65–67^ to the centriole as it converts to a basal body—microtubules and excess PCM around the centriole could act as a direct barrier to the localization of such components or inhibit localization of structures like centriolar satellites that normally facilitate TZ loading. Alternatively, increased microtubule-based transport could perturb the stoichiometry of TZ component recruitment or mistraffic TZ components to other parts of the cell as a result of defects in global neuronal microtubule polarity.

Apart from neuronal morphology defects, another prominent phenotype we observed following CAP-Trap expression was the perdurance of centrioles well past a time when they would normally be eliminated. This protection required multiple PCM components, raising the questions of how PCM can prevent centriole elimination and also how centrioles are normally marked for elimination. Studies from cultured Drosophila DMEL cells similarly support a role for PCM in centriole protection: depletion of PCM led to the precocious loss of centrioles, indicating that components in the cytoplasm actively degrade centrioles once they are exposed ^21^. However, a previous study in *C. elegans* found that removal of a single *C. elegans* PCM protein PCMD-1 did not lead to precocious centriole elimination during oogenesis ^68^, Thus our study supports a mechanism where the complete removal of PCM is necessary to permit centriole elimination.

The mechanisms that mark a naked centriole for degradation are unknown. In Naegleria, centrioles become polyubiquitinated and both the lysosome and proteasome are required during centriole elimination ^69^. Centrioles in *C. elegans* gonads and in Naegleria widen during elimination ^68,69^, and we observed that perduring centrioles appeared to enlarge in CAP-Trap expressing adults relative to earlier larval stages.

Thus, the changes we observed at centrioles likely reflect that CAP-Trap expression delays rather than inhibits the centriole elimination process. In chytrid fungi, axonemal doublet microtubules are degraded once they are reeled into the cytoplasm, but the attached triplet microtubules of the centrioles are left unperturbed ^70^. This centriolar protection in chytrids could be due to a shell of PCM or through a microtubule doublet-, rather than triplet-, specific degradation program. As could be the case in chytrids, the phasmid cytoplasm could be hostile to the microtubule doublets that normally make up the axoneme and temporarily comprise the *C. elegans* centriole just prior to ciliogenesis. Loss of PCM would then normally trigger the degradation of centrioles in this cell type, perhaps through polyubiquitination-based marking for degradation by the proteasome. This model could be extended to the majority of Drosophila cells that normally have centrioles comprised of microtubule doublets where PCM likely protect these doublets until their programmed elimination ^71^. This model still leaves open the question of how organisms with centrioles comprised of triplet microtubules, but which lack PCM like Chlamydomonas cells, trigger the elimination of their centrioles with precise timing, such as at the onset of meiosis ^72,73^. Future studies will be necessary to reveal whether centriole elimination across the tree of life uses similar mechanisms.

While centrosome inactivation is a hallmark of cellular differentiation, studying the consequences of centrosome hyperactivity is difficult, since perturbing the centrosome causes mitotic defects that can prevent cell division and organismal development. Here, we developed CAP-Trap to tissue-specifically and post-mitotically test the function of centrosome inactivation in a differentiated neuron for the first time. Future studies could leverage this tool to determine the consequences of ectopic centrosome activation in other cell types. Neurons might be particularly sensitive to ectopic centrosome activation as they are known to contain relatively few microtubules, especially in axons and dendrites. It will be interesting to determine whether neurons in other organisms are similarly vulnerable to microtubule rewiring by centrosome activation as those in *C. elegans* and whether cells with more elaborate microtubule arrays can be perturbed by centrosome reactivation. For example, cells that comprise myotubes have elaborate microtubule arrays that originate from the nuclear envelope and then form a latticework throughout the cell via Golgi-derived structures ^74,75^. Oocytes similarly develop robust microtubules arrays with large numbers of microtubules built from the cortex ^76^. In these cases where the centrosome might reasonably only generate a small percentage of the overall microtubules in the cell, it is unclear whether centrosomal microtubules would interfere with cellular process as in neurons but will be an exciting test-bed to determine the invasive potential of ectopically active centrosomes. Beyond their role as MTOCs, centrosomes can act as intracellular signaling platforms and cellular signaling is aberrant in many cancers with amplified centrosomes ^77,78^. By harnessing the ability of CAP-Trap to prevent centriole elimination, future studies can test whether the maintenance of centrioles in normally acentriolar differentiated cells perturbs signaling or drives pathogenetic behaviors. Thus, CAP-Trap has numerous interesting future applications and has revealed new functions for microtubules in neurons where centrosome activation leads to severe consequences for neuronal form and function.

## Acknowledgements

We would like to thank Caitlin Taylor, Kang Shen, Pierre Gönczy, and Kevin O’Connell for sharing reagents prepublication. We thank members of the Feldman Lab for helpful feedback and discussions on the project and manuscript. Some of the strains used were from the CGC, which is funded by the NIH Office of Research Infrastructure Programs (P40 OD010440). This work was made possible by support from the NSF GRFP and the CMB Training Grant T32GM007276 awarded to RKN and from R01GM136902, R01GM133950, and R35GM153310 awarded to JLF.

## Author contributions

Conceptualization, R.K.N., J.M., J.L.F., ; methodology, R.K.N., J.M., J.L.F.; formal analysis, R.K.N.; investigation, R.K.N., J.M., and J.L.F.; writing—original draft, R.K.N. and J.L.F.; writing—review and editing, R.K.N., and J.L.F.; visualization, R.K.N., J.M., and J.L.F.; supervision, J.M. and J.L.F.; funding acquisition, R.K.N., J.L.F.

## METHODS

### C. elegans husbandry

*C. elegans* strains were grown Nematode Growth Medium (NGM) plates coated with a lawn of *E. coli* strain OP5082. For all experiments, animals were maintained at room temperature, unless specifically noted.

### Developmental staging of experimental animals

For experiments with adult worms, adult worms were picked off NGM OP50 plates maintained at 20°C , immediately prior to imaging, unless noted. For all experiments with adult animals in Figure 4 and Figure S4A-C, L4 larvae were picked onto NGM plates with OP50, kept at 20°C overnight, and the resulting adult animals were imaged the next day.

For experiments with L2 larvae, gravid adults were bleached (1:2:3 ratio of bleach (Sigma-Aldrich), water, and 1M NaOH) in a microcentrifuge tube, washed 5x in M9, and resulting embryos were kept in a microcentrifuge tube in M9, rotating, overnight at room temperature. Hatched L1 larvae were plated onto NGM plates with OP50 for 22-24 hours at 20°C and then imaged.

For experiments with L1 larvae, gravid adults were spot-bleached onto unseeded plates and kept at 20°C overnight. L1 larvae were picked and prepared for imaging the next day.

For experiments with embryos staged for intestinal E16 stage imaging, gravid adults were incubated in M9 for 4 hours at 20°C , dissected, and the extruded embryos were prepared for imaging.

For experiments with late-stage embryos, 3-fold embryos were picked off NGM OP50 plates containing gravid adults, and the staged embryos were prepared for imaging.

### Allele generation

Endogenous alleles ZF::BFP::SPD-5, TMEM-218::TagRFP, and ZF::PCMD-1 were generated using the CRISPR Self-Excising Cassette (SEC) described preciously (Dickinson et al., 2015). Endogenous alleles mSG::SAS-7 and SPD-2::ZF were generated using the ribonucleoprotein CRISPR method previously described (Ghanta and Mello, 2020). Single-copy transgene maph9p::SAS-4::TagRFP was made using FLP Recombination-Mediated Cassette Exchange (RMCE) as previously described (Nonet, 2020) and inserted in chromosome II using the landing pad strain jsSi1579 within 50 bp of *ttTi5605*.

### Use of Cap-Trap expressing animals

For the majority of experiments, CAP-Trap was expressed from an extrachromosomal array (*wowEx236*). Some experiments, especially those investigating degradation-based suppression of CAP-Trap phenotypes, relied on expression of CAP-Trap from an integrated array (*wowIs4*5 is the integrated array of *wowEx236*). In those cases, the same CAP-Trap array was integrated using the UV/TMP method (detailed protocol below) and the resulting strain was backcrossed 6 times prior to use for experiments in: Figures 1F, 2G-H, 3E-F, 4A-G, 5G-H, 6E-F (SAS-4 experiments only), 6G-J and supplemental figures S3H, S3K, S4B.

### Array integration protocol

Pick 100 L4 worms expressing the array of interest onto non-seeded plates. In the fume hood, prepare a solution of 30ug/mL of 4,5’,8-Trimethylpsoralen, 97.5% (TMP) (Thermofisher) in M9. * TMP is extremely toxic and light sensitive, so work very carefully to avoid spills. Initially dissolve the powder in DMSO before diluting in M9. Ensure that the solution is freshly made (within 6 months). In the fume hood, spread 2 mLs of the TMP solution onto the plate with the array-expressing worms until they are soaked.

Cover the lid of the plate with foil and incubate for 15 minutes. In the Stratalinker, remove the lid from the plate of worms and irradiate worms with 300 uJoules (x100) in the Stratalinker. Place the plate in irradiated worms back in the fume hood with the lid removed to allow for TMP evaporation. Add a drop of OP50 onto the plate to prevent the worms from starving and allow the TMP to evaporate for 1-2 hours. Then, separate the worms onto five OP50 plates (∼20 worms per plate) and allow for the worms to grow until all the food is eaten. Chunk the plate once. To ensure that we isolated integrants as soon as possible, 20 individual F2 animals were singled and designated as individual lines. Those individual lines were allowed to expand and 20 worms from each line was singled. Homozygous integrants were isolated from those plates and backcrossed 6 times before being used for experiments.

### Imaging

#### Preparation of animals for imaging

Prior to imaging, worms were washed in M9, incubated in 2mM levamisole (dissolved in M9) for 10 minutes, and then mounted onto pads made from 5% agarose dissolved in M9. A no. 1.5 coverslip was placed on top and sealed with Vaseline.

For imaging of intestinal E16 stage embryos, Gravid adults were incubated in M9 for 4 hours at 20°C, dissected, and the extruded embryos were mounted onto pads made from 3% agarose dissolved in M9.

Imaging of late-stage embryos was done in an imaging chamber using CO2 immobilization. Embryos were picked off NGM OP50 plates, washed in M9, and then mounted on pads made from 5% agarose dissolved in M9. The pad was inverted onto the coverslip of the imaging chamber. For immobilization, CO2 was flowed through an ozone water humidifier and into the chamber for 5-10 minutes at a rate of 60-80 cc/min until complete immobilization. The chamber was then sealed by closing the entry and exit valve. After the imaging was complete, ambient air was inserted into the chamber using an aquarium air pump through a flow derivation system.

#### Imaging chamber

The imaging chamber for CO2 immobilization was designed using AUTODESK Tinkercad (https://www.tinkercad.com/). The 3D printing files (.stl) are available at NIH 3D 3DPX-022973. The chamber comprises two parts: a bottom part with a square chamber that transforms into a bottom coverslip chamber connected to an exit and entry tubing, surrounded by a moat to house an O-ring; and a top part that seals the chamber’s O-ring. The device was printed on a DLP 3D printer Photon Ultra (Anycubic, Inc.) using a UV curing SLA resin. The printing process involved 3 seconds of on and 1 second of off exposure with a z-step of 50 µm and an XY pixel size of 80 µm. After printing, the two parts were cleaned with 100% isopropanol, and a final cure was performed using a UV blue-light LED for 15 minutes. The device was then dried in an oven for 30 minutes. The top and bottom imaging windows were added and sealed using UV clear resin and 22x40mm #1.5 coverslips. A nitrile rubber O-ring (32x28x2 mm) was added to the chamber’s moat to create a seal. Both the top and bottom parts were lined up with N52 5x2 mm round magnets.

#### Spinning-disk confocal microscopy

Images were acquired on a Nikon Ti-E inverted microscope (Nikon Instruments) equipped with a 1.5x magnifying lens, a Yokogawa X1 confocal spinning disk head, and an Andor Ixon Ultra back thinned EM-CCD camera (Andor), all controlled by NIS Elements software (Nikon). Images were obtained using a 60x Oil Plan Apochromat (NA = 1.4) or 100x Oil Plan Apochromat (NA = 1.45) objective.

#### Image stabilization and correction for videos

For Fig. 1f, the time projection was generated on a single Z-plane EBP-2 movie where bleach correction through histogram matching on ImageJ was used along with denoising using Noise2Void ^80^ software trained on EBP-2 images. Kymographs generated in Fig. 2c’, d’ were made using kymograph clear 2.0A ^81^ on EBP-2 movies that had undergone bleach correction using Fiji and denoising using Noise2Void trained with the same EBP-2 dataset as described above. For Video 3, image stabilization in StackReg, bleach correction in imageJ, and denoising using Noise2Void trained on EBP-2 images was used.

#### Dye filling assay

Gravid adult worms were incubated in 5 μg/mL DiI (1,1′-dioctadecyl-3,3,3′, 3′-tetramethylindocarbocyanine perchlorate, Medchemexpress) diluted in M9 for 1 hour. Worms were then washed 3 times in M9 and placed onto NGM plates for 2 h for destaining. Worms were then mounted and imaged as above.

#### Dye filling for Control ^No C-T^ worms to visualize neuron morphology

Some control animals lacked a marker that would allow for the visualization of phasmid morphology and thus required dye filling to track this feature. In these cases, gravid adult worms were incubated in dye for 5 minutes, washed 5x in M9, incubated in 1mM levamisole for 10 minutes, and then mounted and imaged as above. All Control^No C-T^ worms with the exception of the control animals in Figures 2E, 3H, and 3I were incubated in 250nM diD (Thermo Scientific V22887). Control animals in Figure 2E, 3H, and 3I were incubated in CellBrite blue (Biotium) according to the suggested working stock ((5uL diD + 5uL loading buffer) into 990uL of M9). No dye staining was used for Control^No C-T^ worms in Figures 2G, 3G, 5G, S4A, S4D, or S4G.

### Data analysis

#### EBP-2 kymographs

Kymographs generated in Fig. 2c’, d’ were made using kymograph clear 2.0A ^81^ on EBP-2 movies that had undergone bleach correction using Fiji and denoising using Noise2Void ^80^ trained with a sample EBP-2 dataset. Kymographs in Fig. 3j were generated using the Multi Kymograph plug-in on FIJI without any additional correction. All kymographs were generated on single slice single channel EBP-2 movies. Cellular regions used to generate the kymographs and time steps for each frame are noted in the associated figure legend.

#### Total EBP-2 intensity

EBP-2 Intensity was measured as pixel intensity relative to the background pixel intensity. ROIs were generated in FIJI and used to measure the mean intensity of EBP-2 in the regions of interest (cell body, dendrite middle, dendrite tip) on a summed intensity projection. The same sized ROI was used to measure mean background intensity for correction. The reported enrichment values were calculated as: (mean signal-mean background) / mean background.

#### RAB-11.1 intensity

RAB-11.1 Intensity was measured as pixel intensity relative to the background pixel intensity. For the intensity of RAB-11.1 in the cell body, equal sized ROIs generated in FIJI were used to measure the mean intensity in the cell body and the corresponding background on a summed intensity projection. For the intensity of RAB-11.1 in the dendrite, a segmented line generated in FIJI that was scaled to the thickness of the dendrite was used to measure mean intensity along the length of the dendrite. The reported enrichment values were calculated as: (mean signal-mean background) / mean background.

#### Dendrite Protrusions

Dendrite protrusions were determined as any protrusions from the main dendrite length. Due to the nature of the space-filling morphology marker without a marker for cilia or dendrite identity, in some cases, a severe path finding defect and an ectopic protrusion are indistinguishable and was categorized as an ectopic protrusion for this quantification.

#### Dendrite Length

Dendrite lengths were measured using the segmented line tool in FIJI through Z using the cytoplasmic space-filling marker of the neuron.

#### Dendrite Angle

Dendrite angles were measured using the angle tool in Fiji where the first vector was a straight line down the cell body and the next vector was drawn to the end of the dendrite. The angle reported was the angle between the two vectors—the angle between the cell body and the end of the dendrite.

#### Dendrite length defect categorization

In Figure 4G, dendrites were considered defective in extension if the measured dendrite length was at least two standard deviations shorter than the length of control in Figure 3B.

#### Length Measurements

HMR-1, MAPH-9, and TBB-4 lengths were measured using the segmented line tool on FIJI on summed projection images.

#### HMR-1 intensity

HMR-1 Intensity was measured as pixel intensity relative to the background pixel intensity. Equal sized ROIs generated in FIJI were used to measure the intensity of the posterior-most foci of HMR-1 (likely corresponding to the HMR-1 at the phasmid pore region) on a summed intensity projection. The same sized ROI was used to measure the mean intensity for background correction. The reported enrichment values were calculated as: (mean signal-mean background) / mean background.

#### TMEM-218 intensity

TMEM-218 Intensity was measured as pixel intensity relative to the background pixel intensity. For the intensity of TMEM-218, ROIs generated in FIJI were used on a summed intensity. projection Due to the dim TMEM-218 signal, only the neuron pair closer to the cover slip was used for this analysis. The same sized ROI was used to measure mean background intensity for correction. The reported enrichment values were calculated as: (signal- background) / background.

#### Retracted vs extended dendrite categorization

Dendrites were binned as “retracted” if the measured dendrite length was at least 2 standard deviations below the mean dendrite length of control.

#### SAS-7 foci

SAS-7 foci were determined as enrichments of SAS-7 in the neuron that were at least 50% of the intensity of a centrosome (∼ the intensity of one centriole) in another cell in an equivalent z-slice within the tail region of the same animal. Measurements were done as mean intensity values on a single slice using the same ROI generated in FIJI for the centrosomes used for comparison and the SAS-7 foci measured.

#### SAS-7 intensity measurements

SAS-7 intensity was measured using the same sized ROI generated in FIJI on a single slice for both PCM(+) and PCM(-), C-T expressing animals.

#### MAPH-9 and TBB-4 Intensity Measurements

Intensity was measured on summed intensity projections using ROIs generated in Fiji. The same ROI was used for both control and CAP-Trap neurons.

#### Statistics

For comparisons in which multiple samples were compared, a Kruskal-Wallis test was completed, followed by a Dunn’s multiple comparisons test. For comparisons in which only two samples were compared, a Mann-Whitney-U test was used. Details for the statistical analysis for each comparison used are noted in the figure legends. For all significance thresholds: ns=P>0.05, *=P ≤ 0.05, **=P ≤ 0.01, ***=P ≤ 0.001, ****=P ≤ 0.0001.

---

**Extended Data Figure 1.**
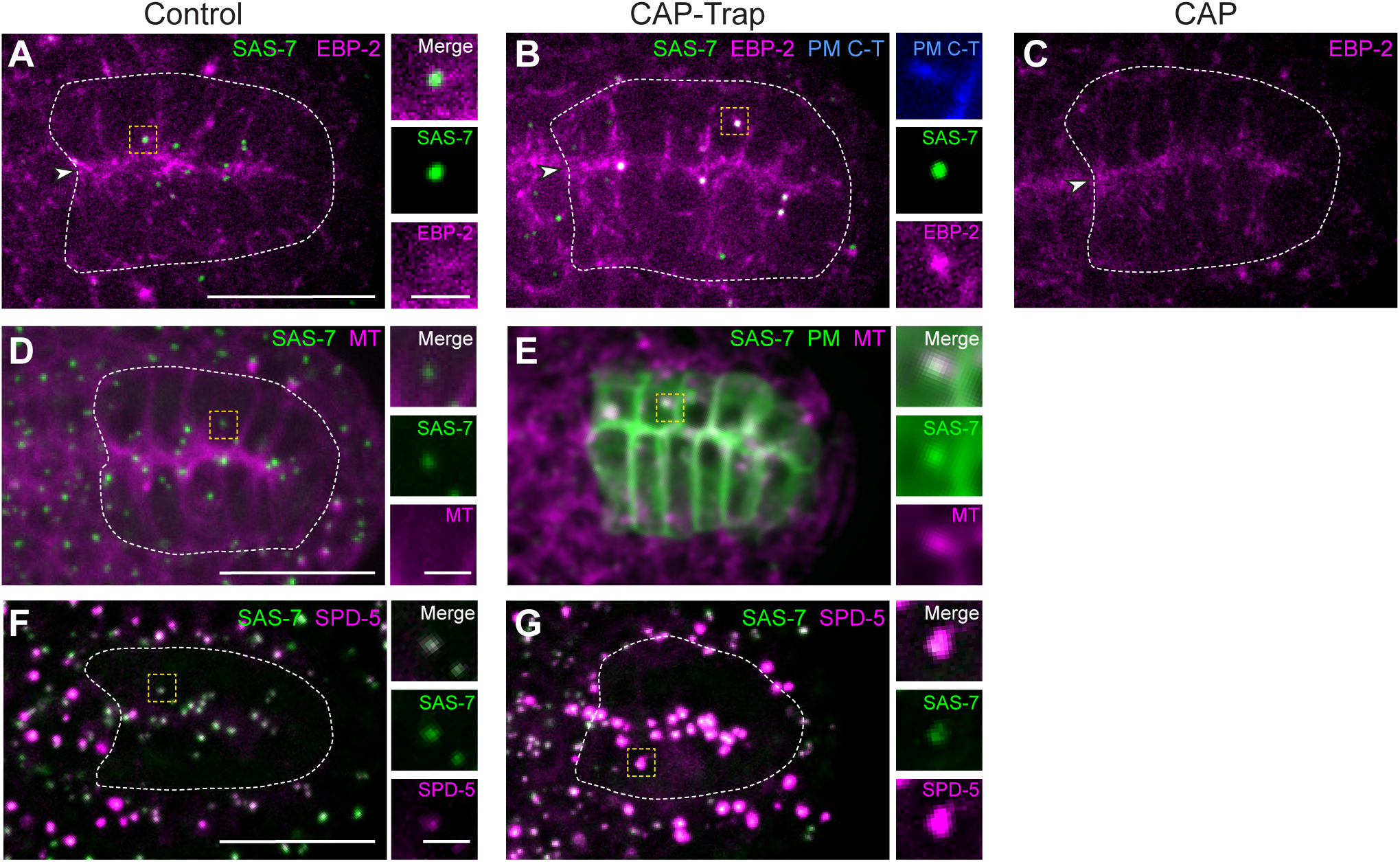
CAP-Trap activates centrosomes in *C*. *elegans* embryonic intestinal cells *C*. *elegans* embryonic intestinal cells at the E16 stage have ncMTOCs at the apical surfaces found at the intestinal midline (arrowhead) that nucleate non-centrosomal microtubules ^2,34^. A-C ) Control^No C-T^ and C-T expressing cells with centrioles (GFP::SAS-7, green throughout) and EBP-2:mKate-2 labeled. Centrioles in Control^No C-T^ embryos do not co-localize EBP-2:mKate-2, indicating that centrosomes are inactive as MTOCs ^2^. CAP-Trap (C-T) expressing cells have centrioles that recruit C-T and EBP-2, indicating that C-T can be successfully targeted to centrioles and ectopically activate MTOC activity at the centrosome. CAP alone control embryos do not express GFP:SAS-7. In both CAP and C-T expressing cell, the plasma membrane marker indicates the expression of the CAP array in the background D-E) Control^No C-T^ and C-T expressing cells with centrioles (GFP::SAS-7) and microtubules (mCherry::TBA-1) labeled. C-T expressing cells also have cell membranes labeled in green (green channel scaled with different LUTs in C-T compared to in control to adjust for bright membrane marker. F-G) Control^No C-T^ and C-T expressing cells with centrioles (GFP::SAS-7) and PCM (TagRFP::SPD-5) labeled. Scale bars = 15μm; Inset scale bars = 2μm. All proteins are endogenously tagged with the exception of mCherry::TBA-1, which is driven by the *pie-1* promoter and inserted into chromosome II. All images are max projections of images acquired using confocal microscopy of live intestinal cells. Magnified insets of boxed regions shown at the right.

**Extended Data Figure 2.**
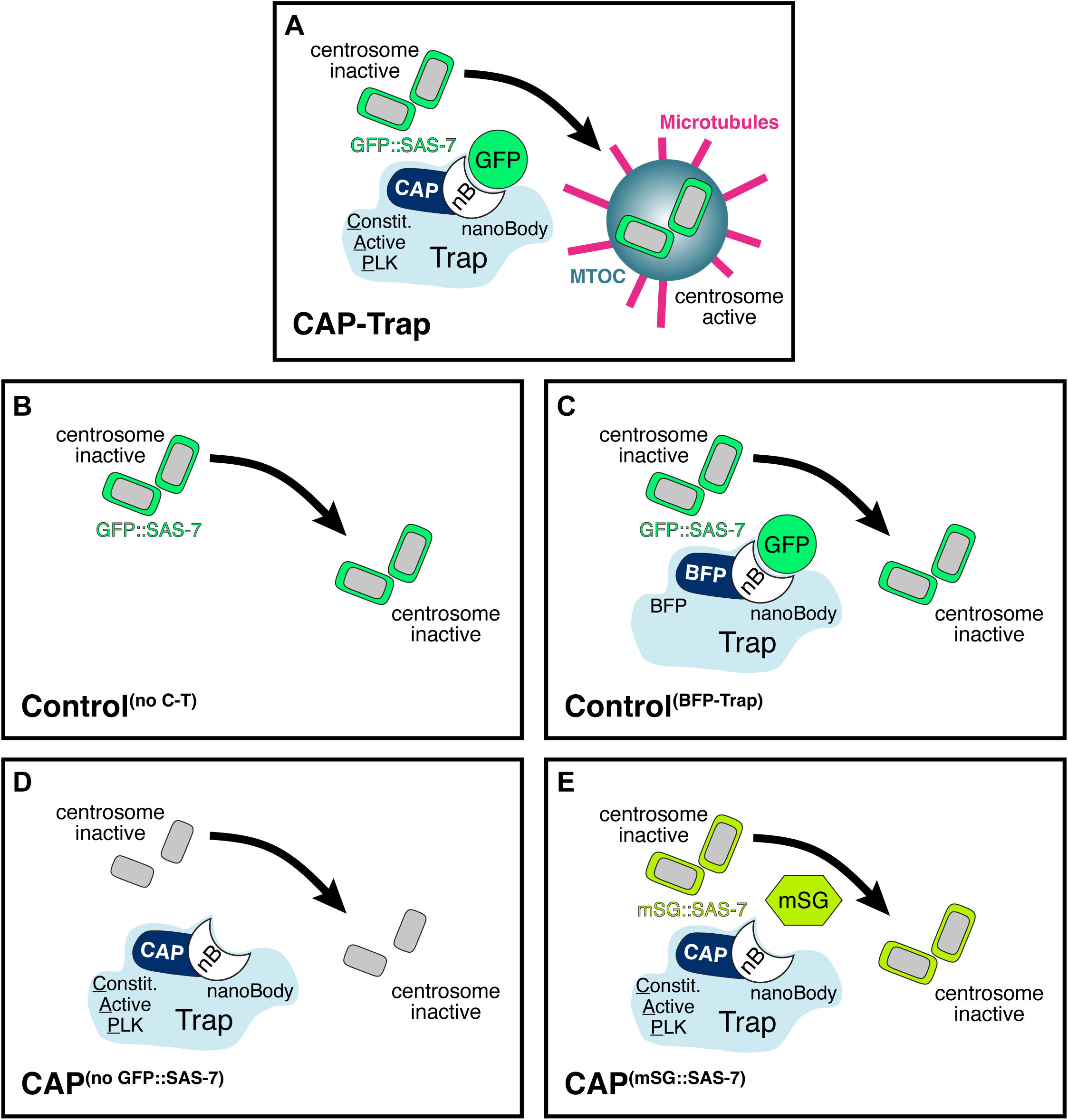
Schematic of control and CAP-Trap tool designs Cartoon schematics of Control^No CT^, Control^BFP-Trap^, CAP, CAP^mSG::SAS-7^, and Cap-Trap experimental designs.

**Extended Data Figure 3.**
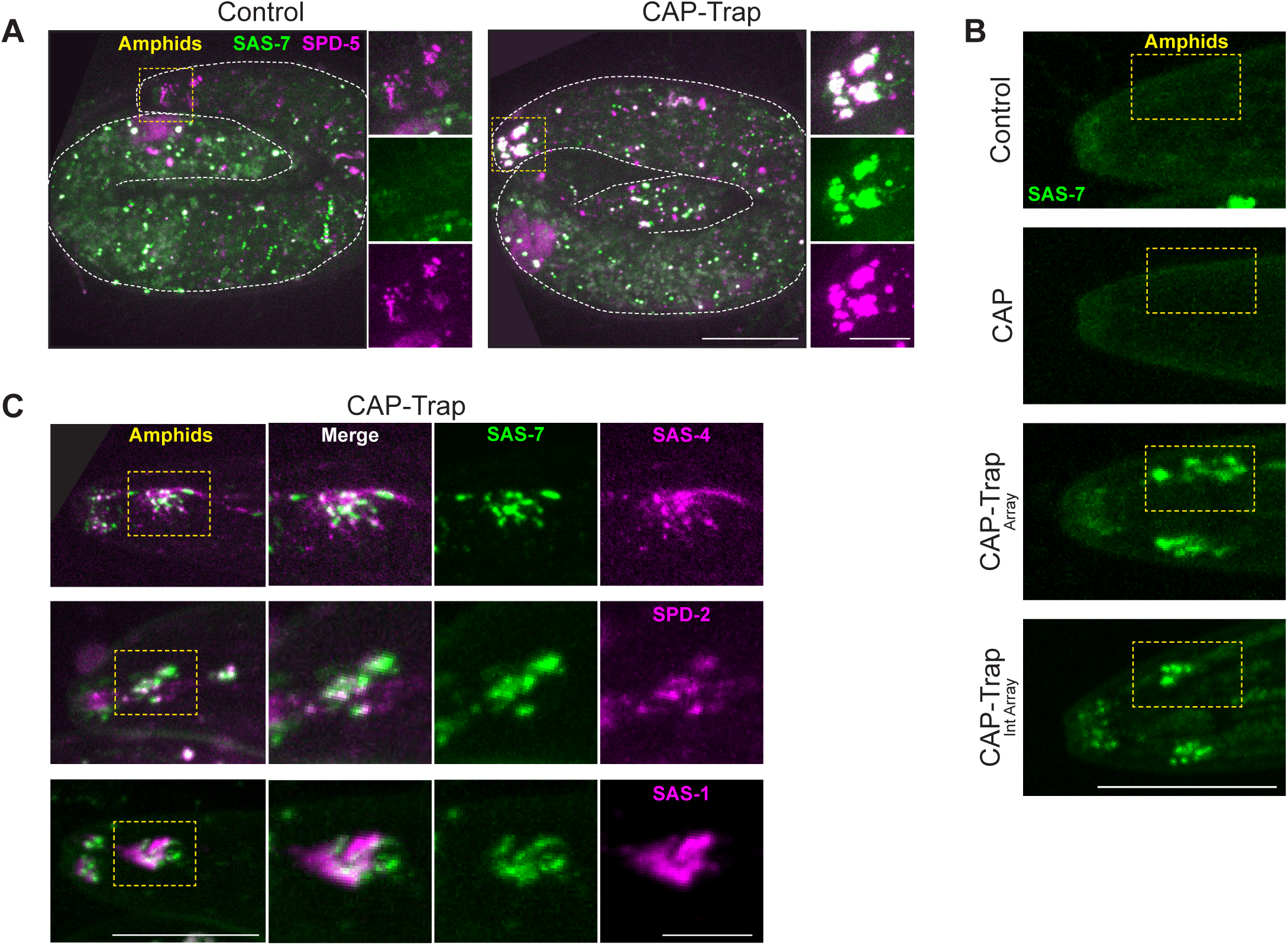
CAP-Trap expression leads to centriole perdurance in the *C*. *elegans* amphid neurons A) Control^No CT^ and C-T^Array^ expressing late-stage 3-fold embryos with centrioles (GFP::SAS-7) and PCM (TagRFP::SPD-5) labeled. Animals were immobilized using a custom imaging chamber (see Methods). Control animals do not have SAS-7 retained due to normal programmed centriole elimination earlier in development but do have SPD-5 localized to the MTOC at the base of each cilium. C-T expressing neurons have centrioles retained with expanded SPD-5. B) Control^No CT^, CAP, C-T^Array^, and C-T^Int Array^ with GFP::SAS-7 labeled in L1 larvae. Centrioles were not retained in Control^No CT^ and CAP expressing neurons while SAS-7 was retained in C-T^Array^ and C-T^Int Array^ expressing neurons. Note that SAS-7 is scaled with the same LUT in Control^No CT^, CAP, and C-T^Array^ conditions but different than C-T^Int Array^ due to differences in imaging parameters. C) Retained GFP::SAS-7 foci in C-T expressing amphid neurons recruit centriolar components SAS-4::TagRFP, TagRFP::SPD-2, and TagRFPT::SAS-1 in L1 larvae. Scale bars =15μm; Inset scale bars = 5μm. All proteins are endogenously tagged with the exception of SAS-4::TagRFP, which is a single-copy insertion driven by the *maph-9* promoter inserted on chromosome II. All images are max projections of images acquired using confocal microscopy of live neurons. Magnified insets of boxed regions shown at the right.

## Supplementary Video Legends

Supplementary Video 1. E16 intestinal cells do not have active centrosomes and microtubules are nucleated from the apical surface Video corresponds to the intestinal primordia in Extended Data Figure 1a. Control^No C-T^ embryonic intestinal cells have centrioles (GFP::SAS-7) and endogenously tagged EBP-2::mKate-2 to mark the apical ncMTOC labeled. Single-plane imaging of EBP-2::mKate-2 dynamics. Centrioles in green are shown only in the first timepoint. Scale bar: 5μm.

Supplementary Video 2. Centrosomes remain inactive in CAP expressing intestinal cells Video corresponds to the intestinal primordia in Extended Data Figure 1b. CAP expressing neurons lack GFP::SAS-7 and express endogenously tagged EBP-2::mKate-2 to mark the apical ncMTOC. Single-plane imaging of EBP-2::mKate-2 dynamics. Scale bar: 5μm.

Supplementary Video 3. CAP-Trap ectopically activates centrosomes in *C. elegans* embryonic intestinal cells Video corresponds to the intestinal primordia in Extended Data Figure 1c. C-T expressing embryonic intestinal cells have centrioles (GFP::SAS-7) and sites of MTOC activity (EBP-2::mKate-2) labeled. Single-plane imaging of EBP-2::mKate-2 dynamics. Centrioles shown in green are only in the first timepoint. Scale bar: 5μm.

Supplementary Video 4. CAP-Trap ectopically activates centrosomes in *C. elegans* ciliated phasmid neurons Video corresponds to the phasmid neuron in Fig. 1F. Phasmid neuron in L2 larvae expressing C-T with centrioles (GFP::SAS-7) and MTOC site (EBP-2::mKate-2) labeled. Individual comet tracks highlighted (arrowhead). Single-plane imaging of EBP-2::mKate-2 dynamics. Scale bar: 5μm.

Supplementary Video 5. EBP-2 dynamics in adult control phasmid neurons Video corresponds to the control phasmid neuron in Fig. 3h. Adult Control^No C-T^ neuron expressing EBP-2::mKate-2 to show sites of microtubule nucleation. Most microtubule dynamics within the neuron are from microtubules nucleated from the MTOC at the base of the cilium at the tip of the dendrite. Inset is of the magnified region of the base of the cilium MTOC. Single-plane imaging of EBP-2::mKate-2 dynamics. Scale bar: 5μm.

Supplementary Video 6. EBP-2 dynamics in CAP-Trap expressing adult phasmid neurons Video corresponds to the phasmid neuron in Fig. 2d. CAP-Trap expressing neuron with EBP-2::mKate-2 showing microtubule nucleation. Plus end out microtubules are nucleated from the centrosome in the cell body and from the ectopic MTOC site in the middle of the dendrite. Inset is of the magnified region of the middle of the dendrite showing mixed microtubule polarity. Single-plane imaging of EBP-2::mKate-2 dynamics. Scale bar: 5μm.

Supplementary Video 7. EBP-2 dynamics in dendrite protrusions in CAP-Trap expressing adult phasmid neurons Video corresponds to the CAP-Trap expressing phasmid neuron in Fig 3h. Video is of the magnified region of the inset in Fig. 3h, showing the ectopic dendrite protrusion. Microtubules are nucleated within the dendrite protrusions. Single-plane imaging of EBP-2::mKate-2 dynamics. Scale bar: 5μm.

## References

1. Sanchez, A. D. & Feldman, J. L. Microtubule-organizing centers: from the centrosome to non-centrosomal sites. Curr Opin Cell Biol 44, 93–101 (2017).

2. Feldman, J. L. & Priess, J. R. A role for the centrosome and PAR-3 in the hand-off of MTOC function during epithelial polarization. Curr Biol 22, 575–582 (2012).

3. Meads, T. & Schroer, T. A. Polarity and nucleation of microtubules in polarized epithelial cells. Cell Motility 32, 273–288 (1995).

4. Mogensen, M. M. Microtubule release and capture in epithelial cells. Biology of the Cell 91, 331–341 (1999).

5. Vergarajauregui, S. et al. AKAP6 orchestrates the nuclear envelope microtubule-organizing center by linking golgi and nucleus via AKAP9. eLife 9, e61669 (2020).

6. Baas, P. W. & Lin, S. Hooks and Comets: The Story of Microtubule Polarity Orientation in the Neuron. Dev Neurobiol 71, 403–418 (2011).

7. Stone, M. C., Roegiers, F. & Rolls, M. M. Microtubules Have Opposite Orientation in Axons and Dendrites of Drosophila Neurons. Mol Biol Cell 19, 4122–4129 (2008).

8. Liang, X. et al. Growth cone-localized microtubule organizing center establishes microtubule orientation in dendrites. eLife 9, e56547 (2020).

9. Ori-McKenney, K. M., Jan, L. Y. & Jan, Y.-N. Golgi outposts shape dendrite morphology by functioning as sites of acentrosomal microtubule nucleation in neurons. Neuron 76, 921–930 (2012).

10. Yagoubat, A. & Conduit, P. T. Asymmetric microtubule nucleation from Golgi stacks promotes opposite microtubule polarity in axons and dendrites. Curr Biol 35, 1311–1325.e4 (2025).

11. del Castillo, U., Winding, M., Lu, W. & Gelfand, V. I. Interplay between kinesin-1 and cortical dynein during axonal outgrowth and microtubule organization in Drosophila neurons. eLife 4, e10140 (2015).

12. Rao, A. N. et al. Cytoplasmic Dynein Transports Axonal Microtubules in a Polarity-Sorting Manner. Cell Rep 19, 2210–2219 (2017).

13. Winding, M., Kelliher, M. T., Lu, W., Wildonger, J. & Gelfand, V. I. Role of kinesin-1–based microtubule sliding in Drosophila nervous system development. Proceedings of the National Academy of Sciences 113, E4985–E4994 (2016).

14. Yan, J. et al. Kinesin-1 regulates dendrite microtubule polarity in Caenorhabditis elegans. eLife 2, e00133 (2013).

15. Zheng, Y. et al. Dynein is required for polarized dendritic transport and uniform microtubule orientation in axons. Nat Cell Biol 10, 1172–1180 (2008).

16. Woodruff, J. B., Wueseke, O. & Hyman, A. A. Pericentriolar material structure and dynamics. Philosophical Transactions of the Royal Society B: Biological Sciences 369, 20130459 (2014).

17. Enos, S. J., Dressler, M., Gomes, B. F., Hyman, A. A. & Woodruff, J. B. Phosphatase PP2A and microtubule-mediated pulling forces disassemble centrosomes during mitotic exit. Biology Open bio.029777 (2017) doi:10.1242/bio.029777.

18. Magescas, J., Zonka, J. C. & Feldman, J. L. A two-step mechanism for the inactivation of microtubule organizing center function at the centrosome. eLife 8, e47867 (2019).

19. Kalbfuss, N. & Gönczy, P. Extensive programmed centriole elimination unveiled in *C. elegans* embryos. Sci. Adv. 9, eadg8682 (2023).

20. Kalbfuss, N. & Gönczy, P. Towards understanding centriole elimination. Open Biology 13, 230222 (2023).

21. Pimenta-Marques, A. et al. A mechanism for the elimination of the female gamete centrosome in Drosophila melanogaster. Science 353, aaf4866 (2016).

22. Zebrowski, D. C. et al. Developmental alterations in centrosome integrity contribute to the post-mitotic state of mammalian cardiomyocytes. eLife 4, e05563 (2015).

23. Godinho, S. A. et al. Oncogene-like induction of cellular invasion from centrosome amplification. Nature 510, 167–171 (2014).

24. Salisbury, J. L., Lingle, W. L., White, R. A., Cordes, L. E. & Barrett, S. Microtubule nucleating capacity of centrosomes in tissue sections. J Histochem Cytochem 47, 1265–1274 (1999).

25. Rusan, N. M. & Wadsworth, P. Centrosome fragments and microtubules are transported asymmetrically away from division plane in anaphase. J Cell Biol 168, 21–28 (2005).

26. McNally, K. L. P. et al. Kinesin-1 prevents capture of the oocyte meiotic spindle by the sperm aster. Dev Cell 22, 788–798 (2012).

27. Erpf, A. C. et al. PCMD-1 Organizes Centrosome Matrix Assembly in C. elegans. Curr Biol 29, 1324–1336.e6 (2019).

28. Hamill, D. R., Severson, A. F., Carter, J. C. & Bowerman, B. Centrosome maturation and mitotic spindle assembly in C. elegans require SPD-5, a protein with multiple coiled-coil domains. Dev Cell 3, 673–684 (2002).

29. Hannak, E. et al. The kinetically dominant assembly pathway for centrosomal asters in Caenorhabditis elegans is gamma-tubulin dependent. J Cell Biol 157, 591–602 (2002).

30. Kemp, C. A., Kopish, K. R., Zipperlen, P., Ahringer, J. & O’Connell, K. F. Centrosome maturation and duplication in C. elegans require the coiled-coil protein SPD-2. Dev Cell 6, 511–523 (2004).

31. Ohta, M. et al. Polo-like kinase 1 independently controls microtubule-nucleating capacity and size of the centrosome. Journal of Cell Biology 220, e202009083 (2021).

32. Woodruff, J. B. et al. Regulated assembly of a supramolecular centrosome scaffold in vitro. Science 348, 808–812 (2015).

33. Tavernier, N. et al. Cdk1 phosphorylates SPAT-1/Bora to trigger PLK-1 activation and drive mitotic entry in C. elegans embryos. J Cell Biol 208, 661–669 (2015).

34. Yang, R. & Feldman, J. L. SPD-2/CEP192 and CDK Are Limiting for Microtubule-Organizing Center Function at the Centrosome. Current Biology 25, 1924–1931 (2015).

35. Wong, S.-S. et al. Centrioles generate a local pulse of Polo/PLK1 activity to initiate mitotic centrosome assembly. EMBO J 41, e110891 (2022).

36. Sulston, J. E., Schierenberg, E., White, J. G. & Thomson, J. N. The embryonic cell lineage of the nematode Caenorhabditis elegans. Dev Biol 100, 64–119 (1983).

37. Magescas, J., Eskinazi, S., Tran, M. V. & Feldman, J. L. Centriole-less pericentriolar material serves as a microtubule organizing center at the base of C. elegans sensory cilia. Curr Biol 31, 2410–2417.e6 (2021).

38. Tran, M. V. et al. MAP9/MAPH-9 supports axonemal microtubule doublets and modulates motor movement. Dev Cell 59, 199–210.e11 (2024).

39. Garbrecht, J., Laos, T., Holzer, E., Dillinger, M. & Dammermann, A. An acentriolar centrosome at the *C. elegans* ciliary base. Current Biology 31, 2418–2428.e8 (2021).

40. Nechipurenko, I. V., Berciu, C., Sengupta, P. & Nicastro, D. Centriolar remodeling underlies basal body maturation during ciliogenesis in Caenorhabditis elegans. Elife 6, e25686 (2017).

41. Serwas, D., Su, T. Y., Roessler, M., Wang, S. & Dammermann, A. Centrioles initiate cilia assembly but are dispensable for maturation and maintenance in C. elegans. J Cell Biol 216, 1659–1671 (2017).

42. Harterink, M. et al. Local microtubule organization promotes cargo transport in C. elegans dendrites. J Cell Sci 131, jcs223107 (2018).

43. Woglar, A. et al. Molecular architecture of the C. elegans centriole. PLOS Biology 20, e3001784 (2022).

44. Armenti, S. T., Lohmer, L. L., Sherwood, D. R. & Nance, J. Repurposing an endogenous degradation system for rapid and targeted depletion of C. elegans proteins. Development 141, 4640–4647 (2014).

45. Sallee, M. D., Zonka, J. C., Skokan, T. D., Raftrey, B. C. & Feldman, J. L. Tissue-specific degradation of essential centrosome components reveals distinct microtubule populations at microtubule organizing centers. PLOS Biology 16, e2005189 (2018).

46. Wang, S. et al. A toolkit for GFP-mediated tissue-specific protein degradation in C. elegans. Development 144, 2694–2701 (2017).

47. Avidor-Reiss, T. Rapid Evolution of Sperm Produces Diverse Centriole Structures that Reveal the Most Rudimentary Structure Needed for Function. Cells 7, (2018).

48. Kobayashi, T. & Dynlacht, B. D. Regulating the transition from centriole to basal body. J Cell Biol 193, 435–444 (2011).

49. Bae, Y.-K. & Barr, M. M. Sensory roles of neuronal cilia: Cilia development, morphogenesis, and function in C. elegans. Front Biosci 13, 5959–5974 (2008).

50. Heiman, M. G. & Shaham, S. DEX-1 and DYF-7 establish sensory dendrite length by anchoring dendritic tips during cell migration. Cell 137, 344–355 (2009).

51. Schouteden, C., Serwas, D., Palfy, M. & Dammermann, A. The ciliary transition zone functions in cell adhesion but is dispensable for axoneme assembly in C. elegans. J Cell Biol 210, 35–44 (2015).

52. Williams, C. L., Winkelbauer, M. E., Schafer, J. C., Michaud, E. J. & Yoder, B. K. Functional redundancy of the B9 proteins and nephrocystins in Caenorhabditis elegans ciliogenesis. Mol Biol Cell 19, 2154–2168 (2008).

53. Williams, C. L. et al. MKS and NPHP modules cooperate to establish basal body/transition zone membrane associations and ciliary gate function during ciliogenesis. J Cell Biol 192, 1023–1041 (2011).

54. Yee, L. E. et al. Conserved Genetic Interactions between Ciliopathy Complexes Cooperatively Support Ciliogenesis and Ciliary Signaling. PLoS Genet 11, e1005627 (2015).

55. Coquand, L. et al. CAMSAPs organize an acentrosomal microtubule network from basal varicosities in radial glial cells. J Cell Biol 220, e202003151 (2021).

56. Thyagarajan, P. et al. Endocytosis of Wnt ligands from surrounding epithelial cells positions microtubule nucleation sites at dendrite branch points. PLOS Biology 23, e3002973 (2025).

57. Fygenson, D. K., Marko, J. F. & Libchaber, A. Mechanics of Microtubule-Based Membrane Extension. Phys. Rev. Lett. 79, 4497–4500 (1997).

58. Ouzounidis, V. R., Prevo, B. & Cheerambathur, D. K. Sculpting the dendritic landscape: Actin, microtubules, and the art of arborization. Current Opinion in Cell Biology 84, 102214 (2023).

59. Pinto-Costa, R. & Sousa, M. M. Microtubules, actin and cytolinkers: how to connect cytoskeletons in the neuronal growth cone. Neuroscience Letters 747, 135693 (2021).

60. Lee, J., Magescas, J., Fetter, R. D., Feldman, J. L. & Shen, K. Inherited apicobasal polarity defines the key features of axon-dendrite polarity in a sensory neuron. Curr Biol 31, 3768–3783.e3 (2021).

61. Stiess, M. & Bradke, F. Neuronal polarization: The cytoskeleton leads the way. Developmental Neurobiology 71, 430–444 (2011).

62. Thyagarajan, P., Feng, C., Lee, D., Shorey, M. & Rolls, M. M. Microtubule polarity is instructive for many aspects of neuronal polarity. Developmental Biology 486, 56–70 (2022).

63. Lindhout, F. W. et al. Centrosome-mediated microtubule remodeling during axon formation in human iPSC-derived neurons. The EMBO Journal 40, e106798 (2021).

64. Jana, S. C. et al. Differential regulation of transition zone and centriole proteins contributes to ciliary base diversity. Nat Cell Biol 20, 928–941 (2018).

65. Gonçalves, J. & Pelletier, L. The Ciliary Transition Zone: Finding the Pieces and Assembling the Gate. Mol Cells 40, 243–253 (2017).

66. Hall, E. A. et al. Centriolar satellites expedite mother centriole remodeling to promote ciliogenesis. eLife 12, e79299 (2023).

67. Klinger, M. et al. The novel centriolar satellite protein SSX2IP targets Cep290 to the ciliary transition zone. MBoC 25, 495–507 (2014).

68. Pierron, M. et al. Centriole elimination during Caenorhabditis elegans oogenesis initiates with loss of the central tube protein SAS-1. EMBO J 42, e115076 (2023).

69. Woglar, A. et al. Mechanisms of axoneme and centriole elimination in Naegleria gruberi. EMBO reports 26, 385–406 (2025).

70. Long, A. F. et al. Dynamic remodeling of centrioles and the microtubule cytoskeleton in the lifecycle of chytrid fungi. MBoC mbc.E24–12-0577 (2025) doi:10.1091/mbc.E24-12-0577.

71. Gottardo, M., Callaini, G. & Riparbelli, M. G. The Drosophila centriole – conversion of doublets into triplets within the stem cell niche. J Cell Sci 128, 2437–2442 (2015).

72. Dutcher, S. K. Elucidation of basal body and centriole functions in Chlamydomonas reinhardtii. Traffic 4, 443–451 (2003).

73. Wingfield, J. L. & Lechtreck, K.-F. Chlamydomonas Basal Bodies as Flagella Organizing Centers. Cells 7, 79 (2018).

74. Musa, H., Orton, C., Morrison, E. E. & Peckham, M. Microtubule assembly in cultured myoblasts and myotubes following nocodazole induced microtubule depolymerisation. J Muscle Res Cell Motil 24, 301–308 (2003).

75. Tassin, A. M., Maro, B. & Bornens, M. Fate of microtubule-organizing centers during myogenesis in vitro. The Journal of cell biology 100, 35–46 (1985).

76. Theurkauf, W. E., Smiley, S., Wong, M. L. & Alberts, B. M. Reorganization of the cytoskeleton during Drosophila oogenesis: implications for axis specification and intercellular transport. Development 115, 923–936 (1992).

77. Arquint, C., Gabryjonczyk, A.-M. & Nigg, E. A. Centrosomes as signalling centres. Philos Trans R Soc Lond B Biol Sci 369, 20130464 (2014).

78. Purkerson, M. M., Amend, S. R. & Pienta, K. J. Bystanders or active players: the role of extra centrosomes as signaling hubs. Cancer Metastasis Rev 44, 1 (2025).

79. Jha, K. et al. C. elegans SAS-1 ensures centriole integrity and ciliary function, and operates with SSNA-1. PLOS Genetics 21, e1011912 (2025).

80. Krull, A., Buchholz, T.-O. & Jug, F. Noise2Void - Learning Denoising From Single Noisy Images. in 2019 *IEEE/CVF Conference on Computer Vision and Pattern Recognition (CVPR)* 2124–2132 (IEEE, Long Beach, CA, USA, 2019). doi:10.1109/CVPR.2019.00223.

81. Mangeol, P., Prevo, B. & Peterman, E. J. G. KymographClear and KymographDirect: two tools for the automated quantitative analysis of molecular and cellular dynamics using kymographs. Mol Biol Cell 27, 1948–1957 (2016).

